# Oceans apart: Heterogeneous patterns of parallel evolution in sticklebacks

**DOI:** 10.1101/826412

**Authors:** Bohao Fang, Petri Kemppainen, Paolo Momigliano, Xueyun Feng, Juha Merilä

**Affiliations:** Ecological Genetics Research Unit, Organismal and Evolutionary Biology Research Program, Faculty of Biological and Environmental Sciences, University of Helsinki, FI-00014 Helsinki, Finland

**Author notes:** These authors contributed equally to this work (Bohao Fang, Petri Kemppainen). Correspondence to: PK and BF.

**Keywords:** *Gasterosteus aculeatus*, genetic differentiation, linkage disequilibrium, local adaptation, parallel evolution

## Abstract

An important model system for the study of genomic mechanisms underlying parallel ecological adaptation in the wild is the three-spined stickleback (*Gasterosteus aculeatus*), which has repeatedly colonized and adapted to freshwater from the sea throughout the northern hemisphere. Previous studies have identified numerous genomic regions showing consistent genetic differentiation between freshwater and marine ecotypes, but these are typically based on limited geographic sampling and are biased towards studies in the Eastern Pacific. We analysed population genomic data from marine and freshwater ecotypes of three-spined sticklebacks with from a comprehensive global collection of marine and freshwater ecotypes to detect loci involved in parallel evolution at different geographic scales. Our findings highlight that most signatures of parallel evolution were unique to the Eastern Pacific. Trans-oceanic marine and freshwater differentiation was only found in a very limited number of genomic regions, including three chromosomal inversions. Using both simulations and empirical data, we demonstrate that this is likely due to both the stochastic loss of freshwater-adapted alleles during founder events during the invasion of the Atlantic basin and selection against freshwater-adapted variants in the sea, both of which have reduced the amount of standing genetic variation available for freshwater adaptation outside the Eastern Pacific region. Moreover, the existence of highly elevated linkage disequilibrium associated with marine-freshwater differentiation in the Eastern Pacific is also consistent with a secondary contact scenario between marine and freshwater populations that have evolved in isolation from each other during past glacial periods. Thus, contrary to what earlier studies focused on Eastern Pacific populations have led us to believe, parallel marine-freshwater differentiation in sticklebacks is far less prevalent and pronounced in all other parts of the species global distribution range.

## Introduction

The extent to which the evolution of similar phenotypes arises by selection acting on shared ancestral polymorphism (i.e. parallel evolution (Schluter & Conte 2009)) or via distinct molecular evolutionary pathways (i.e. convergent evolution (Arendt & Reznick 2008; DeFaveri *et al*. 2011; Stern 2013) is a major question in evolutionary biology. A powerful approach to disentangle these processes is to study the genomic architecture underlying the repeated evolution of similar phenotypes in similar environments. After the retreat of Pleistocene glaciers, marine three-spined sticklebacks (*Gasterosteus aculeatus*) colonized and adapted to many newly formed freshwater habitats by adopting similar changes in a number of morphological, physiological, life history and behavioural traits (Bell & Foster 1994; Gibson 2005; Hendry *et al*. 2013; Lescak *et al*. 2015; Östlund-Nilsson *et al*. 2006). Thus, this species has become one of the most widely used model systems to study the molecular basis of adaptive evolution in vertebrates in the wild (McKinnon & Rundle 2002).

Previous studies of the three-spined stickleback model system have quantified the extent of parallel evolution by identifying genomic regions that are consistently differentiated between marine and freshwater ecotypes sampled across different geographic areas (DeFaveri *et al*. 2011; Ferchaud & Hansen 2016; Hohenlohe *et al*. 2010; Hohenlohe & Magalhaes 2019; Jones *et al*. 2012; Liu *et al*. 2018; Pujolar *et al*. 2017; Terekhanova *et al*. 2019; Terekhanova *et al*. 2014). The focus has historically been on the Eastern Pacific region (Chan *et al*. 2010; Colosimo *et al*. 2005; Hohenlohe *et al*. 2010; Hohenlohe & Magalhaes 2019; Jones *et al*. 2012; Nelson & Cresko 2018), but several recent studies have focused on Atlantic populations (Ferchaud & Hansen 2016; Liu *et al*. 2018; Pujolar *et al*. 2017; Terekhanova *et al*. 2019; Terekhanova *et al*. 2014). However, only two studies have thus far included samples from a larger (global) geographic range (DeFaveri *et al*. 2011; Jones *et al*. 2012). Based on whole genome sequence data from a limited number of individuals from Eastern Pacific and Atlantic populations (n = 21), Jones *et al*. (2012) identified ~200 genomic regions that consistently separated marine and freshwater individuals globally, representing roughly 0.5% of the dataset. They also found that 2.83% of the genome showed signatures of parallel selection in Eastern Pacific freshwater locations – approximately six times more than that at the global scale (tree i, Supplementary Fig. 2 and Supplementary Table 7 in Jones *et al*. (2012) – suggesting that more loci contribute to parallel evolution at smaller geographic (regional) scales. However, since this pattern was unique to the Eastern Pacific (and the focus was on global parallelism) its implications for a holistic understanding of marine-freshwater differentiation at both regional and global scales was never discussed. Such global heterogeneous ecotype divergence is consistent with the results of several other studies as well. Focusing on 26 candidate genes in six pairs of marine-freshwater populations across the globe, DeFaveri *et al*. (2011) found that only ~50% of the genes under divergent selection were shared across more than three population pairs, and none were shared among all populations. This suggested a limited re-reuse of ancestral polymorphism at the global scale, implicating either an important role of convergent evolution at larger geographic scales (DeFaveri *et al*. 2011), or geographic heterogeneity in selective pressure among different (DeFaveri *et al*. 2011) freshwater ecosystems (DeFaveri *et al*. 2011; Stern 2013). Furthermore, studies focusing on parallel evolution within oceans, and even smaller geographic regions, show striking differences in the proportion of loci involved in parallel freshwater adaptation between Pacific and Atlantic regions (Ferchaud & Hansen 2016; Hohenlohe *et al*. 2010; Jones *et al*. 2012; Liu *et al*. 2018; Nelson & Cresko 2018; Pujolar *et al*. 2017; Terekhanova *et al*. 2019; Terekhanova *et al*. 2014). For instance, Terekhanova *et al*. (2014, 2019) recovered only 21 highly localized genomic regions involved in parallel freshwater differentiation in the White Sea, in contrast to Hohenlohe et al. (2010) and Nelson & Cresko (2018) who found large genomic regions involved in parallel freshwater differentiation across almost all chromosomes in the Eastern Pacific populations. Therefore, the potential mechanisms underlying this apparent large-scale geographic heterogeneity in genome-wide patterns of parallel evolution in three-spined sticklebacks remain unexplored. To this end, we analysed population genomic data from a comprehensive sampling of all major geographic areas inhabited by the three-spined stickleback, and employed unsupervised and supervised methods to detect loci involved in parallel marine-freshwater differentiation at different geographical scales. Based on earlier observations (DeFaveri *et al*. 2011; Ferchaud & Hansen 2016; Kemppainen *et al*. 2015; Liu *et al*. 2018; Pujolar *et al*. 2017; Terekhanova *et al*. 2019; Terekhanova *et al*. 2014), we hypothesize that the genetic parallelism in response to freshwater colonization by marine sticklebacks is heterogeneous at the global scale, and that the degree of genetic parallelism is much stronger in the Eastern Pacific region than elsewhere.

We further seek to understand and discuss the ultimate causes of the marked regional differences in genome-wide signatures of parallel genetic differentiation among ecotypes. To explain the mechanism behind the repeated use of the same alleles in independent freshwater populations of sticklebacks, Schluter & Conte (2009) proposed the “transporter hypothesis”. This hypothesis postulates that three-spined sticklebacks have repeatedly colonized and adapted to freshwater environments via selection on standing genetic variation in large marine populations. These freshwater-adapted alleles are in turn maintained in the marine populations by recurrent gene flow with previously colonized freshwater populations. Three-spined stickleback populations have persisted in the Eastern Pacific for approximately 26 Mya (Betancur *et al*. 2015; Matschiner *et al*. 2011;

Meynard *et al*. 2012; Sanciangco *et al*. 2016) and from there recolonized the Western Pacific and Atlantic Ocean basins following local extinctions much more recently, during the late Pleistocene (36.9-346.5 kya; Fang *et al*. 2020; Fang *et al*. 2018; Orti *et al*. 1994). During bottlenecks and founder events, rare alleles are lost at a higher rate than common alleles (Halliburton & Halliburton 2004; Hyten *et al*. 2006). Since freshwater-adapted alleles exist in the marine populations only at low frequencies (Schluter & Conte 2009), it is likely that they were lost to a higher degree than neutral variation during geographic range expansions from the Eastern Pacific (via the sea), thereby reducing the amount of standing genetic variation available for freshwater adaptation outside of the Eastern Pacific. To test this hypothesis, we used individual-based forward simulations designed to mimic the transporter hypothesis, and the general global population demographic history of three-spined sticklebacks outlined above. We conclude with a discussion on other potential biological and demographic explanations for the high degree of geographic heterogeneity in patterns of parallel genomic differentiation, and reflect upon the representativeness of the Eastern Pacific three-spined stickleback populations as a general model for the study of parallel evolution.

## Material and Methods

### Sample collection

We obtained population genomic data from 166 individuals representing both marine and freshwater ecotypes from the Eastern and Western Pacific, as well as from the Eastern and Western Atlantic Oceans (Fig. 1i, Supplementary Table 1 and Supplementary Fig. 1). Additional data from previously published studies were retrieved from GenBank. Fish collected for this study were sampled with seine nets, minnow traps and electrofishing. Specimens were preserved in ethanol after being euthanized with an overdose of Tricaine mesylate (MS222).

**Figure 1.**
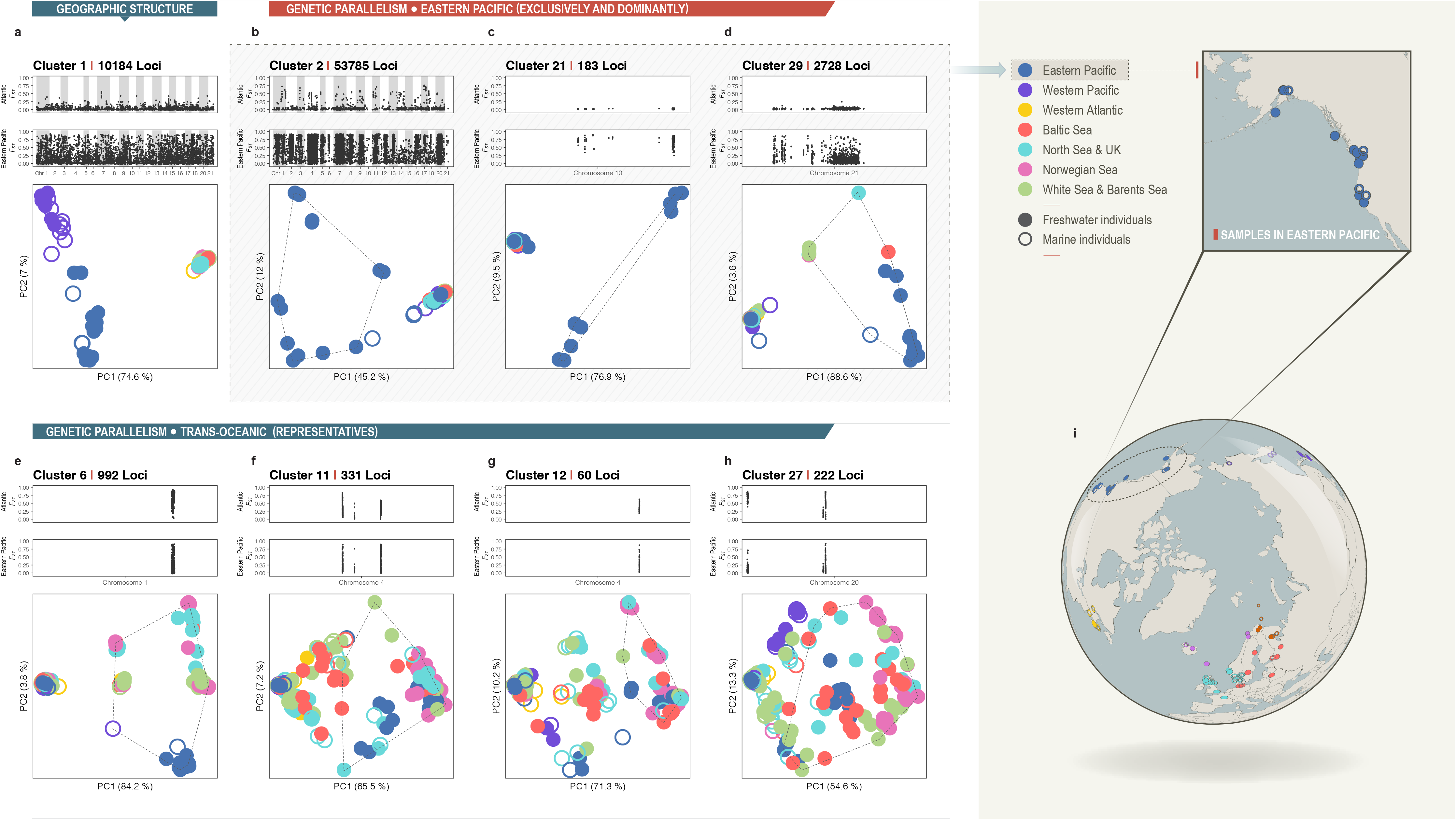
Linkage Disequilibrium network analysis (LDna). (a-h) Eight main clusters of loci identified by LDna (LD-clusters). In each panel (LD-cluster), the top and middle plots present the marine-freshwater differentiation (*F*_ST_) between Atlantic and Eastern Pacific samples, respectively. The bottom plot shows the principal component analysis (PCA) based on the LD-cluster loci. The seven different colours represent the geographic origin of populations. Solid and open circles refer to freshwater and marine ecotypes, respectively. All identified LD-clusters (29 in total) and corresponding information are presented in Supplementary Fig. 2 and Supplementary Table 2. (j) Map of the sampled populations; colours match those in the PCA results. A Mercator projection of the sampling map is shown in Supplementary Fig. 1.

To study the extent of genetic parallelism among freshwater sticklebacks with different phylogeographic histories, we classified global samples into seven biogeographic regions based on their phylogenetic affinities: (i) Eastern Pacific, (ii) Western Pacific, (iii) Western Atlantic, (iv) White and Barents Seas, (v) North Sea and British Isles, (vi) Baltic Sea and (vii) Norwegian Sea (Fang et al., 2018; Fig. 1i). A summary of coordinates, ecotype and population information on the sampled individuals and re-acquired samples is given in the Supplementary Table 1.

### Sequencing and genotype likelihood estimation

Restriction site associated DNA sequencing (RADseq) using the enzyme *PstI* was performed for the 62 individuals sampled in this study, using the same protocol as in Fang et al. (2018), where DNA library preparation and sequencing method are described in detail. The raw RAD sequencing data has been uploaded to GenBank (Accessions SAMN14078677-SAMN14078738). Previously published RADseq and whole genome sequencing (WGS) data for an additional 104 individuals from 62 populations were retrieved from GenBank (Supplementary Table 1). All RADseq and WGS datasets were mapped to the three-spined stickleback reference genome (release-92, retrieved from Ensembl (Hubbard *et al*. 2005) using BWA mem v0.7.17 (Li & Durbin 2009). PCR duplicates were removed using the program Stacks v2.5 (Catchen *et al*. 2013) for pair-end RAD data, and SAMtools v1.9 (function “markdup”(Li 2011)) for whole genome data. Given the heterogeneity in sequencing depth among different datasets, and particularly the very low coverage of the data retrieved from Jones et al. (Jones *et al*. 2012), most of the analyses were performed directly using genotype likelihoods, avoiding variant calling whenever possible. Genotype likelihoods where estimated from the mapped reads using the model of SAMtools (Li 2011) as implemented in the program suite ANGSD v0.929 (Korneliussen *et al*. 2014). Full scripts for the genotyping and filtering parameters are publically available through DRYAD (doi: xxx). Bases with a q-score below 20 (-minQ 20) and reads with mapping quality below 25 (-minMapQ 25) were removed, and variants were only retained if they had a p-value smaller than 1e-6 (-SNP_pval 1e-6 flag in ANGSD). We retained sites with a minimum read depth of two (-minIndDepth 2) in at least 80% of the sampled individuals (-minInd 133). The sex chromosome (Chr. XIX (Kitano *et al*. 2009; Natri *et al*. 2013)) was excluded from downstream analyses due to sex-specific genomic heterogeneity (Hedrick 2007; Schaffner 2004). The raw output of genotype likelihoods from all 166 individuals comprised 2,511,922 genome-wide loci.

### Unsupervised approach to determine marine-freshwater differentiation

We conducted Linkage Disequilibrium Network Analysis (LDna) on the whole dataset (2,511,922 SNPs) to identify and extract clusters of highly correlated loci, i.e. sets of loci affected by the same evolutionary processes. LDna uses a pairwise matrix of LD values, estimated by *r_2_*, to produce a single linkage clustering tree. The hierarchical clustering algorithm uses the LD matrix to combine two clusters connected to each other by at least one edge. In the resulting tree, the nodes represent clusters of loci connected by LD values above thresholds, where the threshold value is proportional to the distance from the root (Kemppainen *et al*. 2015). As the LD threshold is sequentially lowered, an increasing number of loci will be connected to each other in a fashion that reflects their similarity in phylogenetic signals. For each cluster merger (with decreasing LD threshold), the change in median LD between all pairwise loci in a cluster before and after the merger is estimated as λ. When two highly interconnected clusters merge, λ will be large (unlike when only a single locus is added to an existing cluster), signifying that these two clusters bear distinct phylogenetic signals. LDna is currently limited to ~20,000 SNPs at a time due to its dependence on LD estimates for all pairwise comparisons between loci in the dataset. To analyse the whole dataset, we applied a novel three-step LDna approach to reduce the complexity of the data in a nested fashion. First, we started with non-overlapping windows within each chromosome (Li *et al*. 2018), then performed the analysis on each chromosome individually, and finally on the whole dataset (Supplementary Information 1).

In all steps of LDna, we estimated LD between loci from genotype likelihoods using the program ngsLD(Fox *et al*. 2019), setting the minimum SNP minor allele frequency at 0.05. Full scripts for the LDna analyses are provided in DRYAD (doi: xxx). In the first step, we only kept loci that were in high LD with at least one locus (*r_2_*>0.8) within a window of 100 SNPs, as most SNPs in the data were not correlated with any other adjacent loci (so-called singleton clusters (Li *et al*. 2018), and thus, are unlikely to be informative in the LDna analyses.

The main evolutionary phenomena that cause elevated LD between large sets of loci in population genomic datasets are polymorphic inversions, population structure and local adaptation, all of which are expected to be present in our dataset (Kemppainen *et al*. 2015). There are specific and distinct predictions about the population genetic signal and the distribution of loci in the genome that arise from these evolutionary phenomena (Kemppainen *et al*. 2015). First, clusters with LD signals caused by inversions are expected to predominantly map to the specific genomic region where the inversion is situated. In addition, a Principal Component Analysis (PCA) of these loci is expected to separate individuals based on karyotype. In general, the heterokaryotype would be intermediate to the two alternative homokaryotypes (provided that all karyotypes exist in the dataset), and the heterokaryotypic individuals would show higher observed heterozygosity than the homokaryotypes. However, this is not always so clear, for instance when the inversion is new and mutational differences have not yet accumulated. Second, a PCA based on loci whose frequencies are shaped by genetic drift is expected to separate individuals on the basis of geographic location, with no (or very little) separation between marine and freshwater ecotypes. Third, an LD signal caused by local adaptation (globally) is expected to cluster individuals based on ecotype, regardless of geographic location, with both the locus distribution and LD patterns being negatively correlated with local recombination rate to some extent (Roesti *et al*. 2014; Roesti *et al*. 2013). The reason for this correlation is that gene flow between ecotypes erodes genetic differentiation in sites linked to locally adapted loci with the exception of regions where recombination is restricted (for instance in inversions, or close to centromeres or telomeres). No such pattern is expected for LD caused by population structuring, as the main source of this LD is the random genetic drift that, in the absence of gene flow, generates LD in a fashion that is independent of genome position (Kemppainen *et al*. 2015) and background selection is not expected to result in strong patterns of genetic differentiation across genomes (Matthey-Doret & Whitlock 2019; Stankowski *et al*. 2019). If a set of loci contributes to local adaptation exclusively in a particular geographic area, a PCA based on these loci will only separate individuals based on ecotype in that region. We considered loci to be involved in parallel evolution only if they grouped individuals of the same ecotype from more than one independent location. Otherwise, it is not possible to discern drift from local adaptation, particularly if *N_e_* is small (i.e. genetic drift is strong). To determine if an LD-cluster was likely associated with parallel freshwater differentiation, we first used expectation maximisation (EM) and hierarchical clustering methods to identify clusters of individuals in PCAs that contained a minimum of seven individuals, of which at least 90% are freshwater ecotypes (the “in-group”; dotted line; Fig. 1a-h, Supplementary Fig. 2). Second, if such in-groups were detected, we used permutations to further test whether this cluster contained more freshwater individuals than expected by chance (Supplementary Information 2). With less than seven in-group individuals, there was no power to detect significant associations, even if all individuals were freshwater ecotypes. We benchmarked LDna by quantifying the proportion of regions previously identified by Jones et al. (2012) as involved in marine-freshwater ecotype differentiation (globally and within the Eastern Pacific) that were correctly recovered by LD-cluster loci (Supplementary Information 5).

### Supervised approaches to determine marine-freshwater differentiation

Genome-wide allelic differentiation (*F*_ST_ estimated from genotype likelihoods in ANGSD) between marine and freshwater ecotypes was estimated separately for the three major oceans in our study: Eastern Pacific, Western Pacific and Atlantic Oceans. All available samples were always used, but due to the small number of available Pacific marine individuals, marine individuals from the Eastern (n=4) and Western Pacific (n=13) were pooled and treated as a combined Pacific marine group in Eastern and Western Pacific ecotype comparisons (Supplementary Table 2) in order to reduce the noise of the SNP-based analyses. To determine whether pooling Easter and Western Pacific marine individuals could bias *F*_ST_ estimates, we first estimated *F*_ST_ in 100 kb windows for Eastern Pacific marine *vs*. Eastern Pacific freshwater and Western Pacific marine *vs*. Eastern Pacific freshwater individuals, as using large windows allowed us to get precise estimates even if we sampled only four marine individuals from the Eastern Pacific. The two sets of window-based pairwise *F*_ST_ estimates were highly correlated (*r* = 0.904; P<0.001; Supplementary Fig. 4d-g), suggesting that pooling marine individuals from Eastern and Western Pacific should not strongly affect SNP based estimates. Note that from the results of the unsupervised LDna, two Eastern Pacific freshwater individuals from Kodiak Island, Alaska (ALA population) never grouped with the other Eastern Pacific freshwater individuals. Therefore, in agreement with earlier phylogenetic analyses (Fang *et al*. 2018), these two individuals were excluded from the supervised analyses. The squared correlation coefficient of *F*_ST_ before and after this exclusion was 0.88, indicating that this did not affect the results.

In each comparison, the sites were firstly filtered from raw mapped reads, retaining sites with less than 25% missing data with quality control (-minIndDepth 1, -uniqueOnly 1, -remove_bads 1, -minMapQ 20, -minQ 20). We retained only variable sites (-SNP_pval 1e-6) in each region, resulting in 1,218,858 SNPs in the Eastern Pacific, 1,072,257 SNPs in the Western Pacific, and 1,681,923 SNPs in the Atlantic Ocean. We then obtained genotype likelihoods and site allele frequency likelihoods of the variants (-GL 1, -doSaf 1). Based on these likelihoods, we estimated the two-dimensional site-frequency spectrum (SFS) for each pair of ecotypes (realSFS) and calculated the pairwise weighted *F*_ST_ (realSFS fst).

### Proof of concept using simulated data

Several potential explanations for geographic heterogeneity in parallel patterns of marine-freshwater differentiation in three-spined sticklebacks have been suggested (DeFaveri *et al*. 2011). One such explanation that has not received much attention in the context of three-spined sticklebacks is the stochastic loss of freshwater-adapted alleles due to founder events when three-spined sticklebacks colonized the rest of the world from the Eastern Pacific in the late Pleistocene (see Introduction). Thus, as a proof of concept, we used forward-in-time simulations performed with quantiNemo (Neuenschwander *et al*. 2008) to investigate the conditions under which parallel islands of differentiation between marine and freshwater ecotypes can arise under such a scenario.

Our simulations were aimed at recreating the transporter hypothesis model in the Eastern Pacific (referred to as *“Pac”* in the context of simulations), to simulate the colonization of the Atlantic (referred to as *“Atl”* in the context of simulations) from *Pac* 60-30 Kya during the last known opening of the Bering Strait(Fang *et al*. 2020; Hu *et al*. 2010; Meiri *et al*. 2014) and the subsequent post-glacial (10 Kya) colonization of newly formed freshwater habitats in both oceans (simulation details can be found in Supplementary Information 4). In short, simulations begin with one marine population in *Pac* connected to five independent freshwater populations by symmetrical gene flow (i.e. no gene flow exists between any of the freshwater populations; Fig. 3a) for 10k generations (40-50 kya). This is followed by colonisation of *Atl* from *Pac* (with *Atl* having identical population structure to *Pac*) by allowing one or five migrants per generation between the oceans for 2 Ky (38-40 Kya), after which no further gene flow is possible. The retreat of the Pleistocene continental ice sheets (at 10 Kya; Fig. 3d) and the colonization of newly formed freshwater habitats is simulated by removing four of the freshwater populations, immediately followed by the emergence of four new (post-glacial) freshwater populations (in both *Pac* and *Atl*; Fig. 3e). The fifth freshwater population remains as a “glacial refugia” that continues to feed freshwater-adapted alleles to the sea as standing genetic variation. Post-glacial local adaptation is thus only possible due to the spread of freshwater-adapted alleles from the sea in accordance with the transporter hypothesis (Schluter & Conte, 2009; Fig. 3a-e).

Marine-freshwater differentiation was based on bi-allelic QTL with allelic effects of either zero or ten, with the selection optima in the marine habitat being zero and the selection optima in all freshwater populations being 20. Thus, a freshwater individual homozygous for allele 2 for a given QTL meant that the individual was at its optimal phenotype, and *vice versa* for marine individuals. Selection intensities were such that a sufficient amount of standing genetic variation was allowed in the sea and rapid local adaptation in freshwater was possible (see Supporting Information 4 for details). In simulations, all allele frequencies started from 0.5 in all populations (including the QTL in the freshwater habitats). The simulated genome was comprised of ten equally sized chromosomes, with a total genome size of 1000 bps. Regions of both low (centromeric regions) and high (chromosome arms) recombination were represented (Supplementary Information 4). Either 3, 6 or 9 QTL per chromosome were randomly placed in eight of the chromosomes, after which the positions were fixed, leaving the last two chromosomes without any QTL. Twenty replicate simulations were run for each of the six different parameter settings (two levels of trans-oceanic gene flow rates and three different QTL densities). The frequency of freshwater-adapted alleles was recorded at 50-generation intervals throughout the simulations. Population genomic data were saved at the end of the simulations (representing present-day sampling).

### Linking empirical data to simulated data

LDna identified one major cluster (LD-cluster 2; see Results) that separated all Eastern Pacific freshwater individuals from the remaining individuals (Atlantic, Western Pacific and marine samples from the Eastern Pacific, pooled). From the simulated data, we first sub-sampled individuals from *Pac* and *Atl* to match the samples size of the empirical data (excluding the Western Pacific samples, as this ocean was not included in the simulations), and used LDna to detect clusters similar to LD-cluster 2 using Cluster Separation Scores (CSS; custom R-scripts available from DRYAD). Cluster Separation Scores were calculated as the Euclidean centroid distance in a PCA (based on coordinates from the two first principal components scaled by their eigenvalues) between two groups of individuals, standardized by the longest distance between any two individuals in the PCA (CSS thus ranging between [0,1]). PCA of the simulated datasets were performed by the function snpgdsPCA from the R-package SNPRelate(Zheng *et al*. 2012). CSS scores are known to correlate with *F*_ST_, but give higher resolution when genetic differentiation is high, and are less sensitive to small sample sizes(Jones *et al*. 2012). In LDna, we are only interested in clusters with high λ-values (see above and Supplementary Information 1). Therefore, from the ten LD-clusters with the highest λ-values (from each simulated dataset), we considered the cluster with the highest CSS between *PF* (Eastern Pacific freshwater) and *non-PF* individuals to be the strongest candidates for showing high ecotype differentiation specifically in *Pac*, and thus, the most similar to LD-cluster 2 in the empirical data. The non-*PF* individuals were comprised of *PM* (Eastern Pacific marine), *AF* (Atlantic freshwater) and *AM* (Atlantic marine) individuals pooled. To further assess how similar the patterns of population differentiation (in PCAs) were in the above LD-clusters (simulated data) and empirically obtained LD-cluster 2, we compared the CSS’s for all pairwise comparisons between *PF* and the other three groups of individuals (i.e. “*PF vs. PM”, “PF vs. AF”* and “*PF vs. AM”*) in the simulated and empirical datasets. To further assess the extent to which clusters similar to LD-cluster 2 could be produced in the *Atl*, we used the same procedure as above to look for the LD-clusters with the highest CSS scores between *AF* and non-*AF* individuals (*AM*, *PF* and *PM*), and calculated CSS scores between *AF* individuals and the three non-*AF* groups.

## Results

### Marine-freshwater divergence determined by unsupervised and supervised approaches

The first step of LDna on the empirical dataset (2,511,922 SNPs derived from 166 individuals worldwide) identified 214,326 loci that were in high LD with at least one other locus within windows of 100 SNPs (Supplementary Information 1). The next step of performing LDna on each chromosome separately (only using one locus from each LD-cluster from step one; Supplementary information 1) resulted in 81 distinct LD-clusters. From these, a final 29 LD-clusters were obtained (pooling within chromosome LD-clusters whenever they were grouped by LDna in the final step; Supplementary information 1), containing a total of 71,064 loci (*viz*. Cluster 1-29). Eight of these LD-clusters associated with geographic structure and genetic parallelism are highlighted in Fig. 2a-h. Details of all 29 clusters can be found in Supplementary Fig. 2 and Supplementary Table 3.

**Figure 2.**
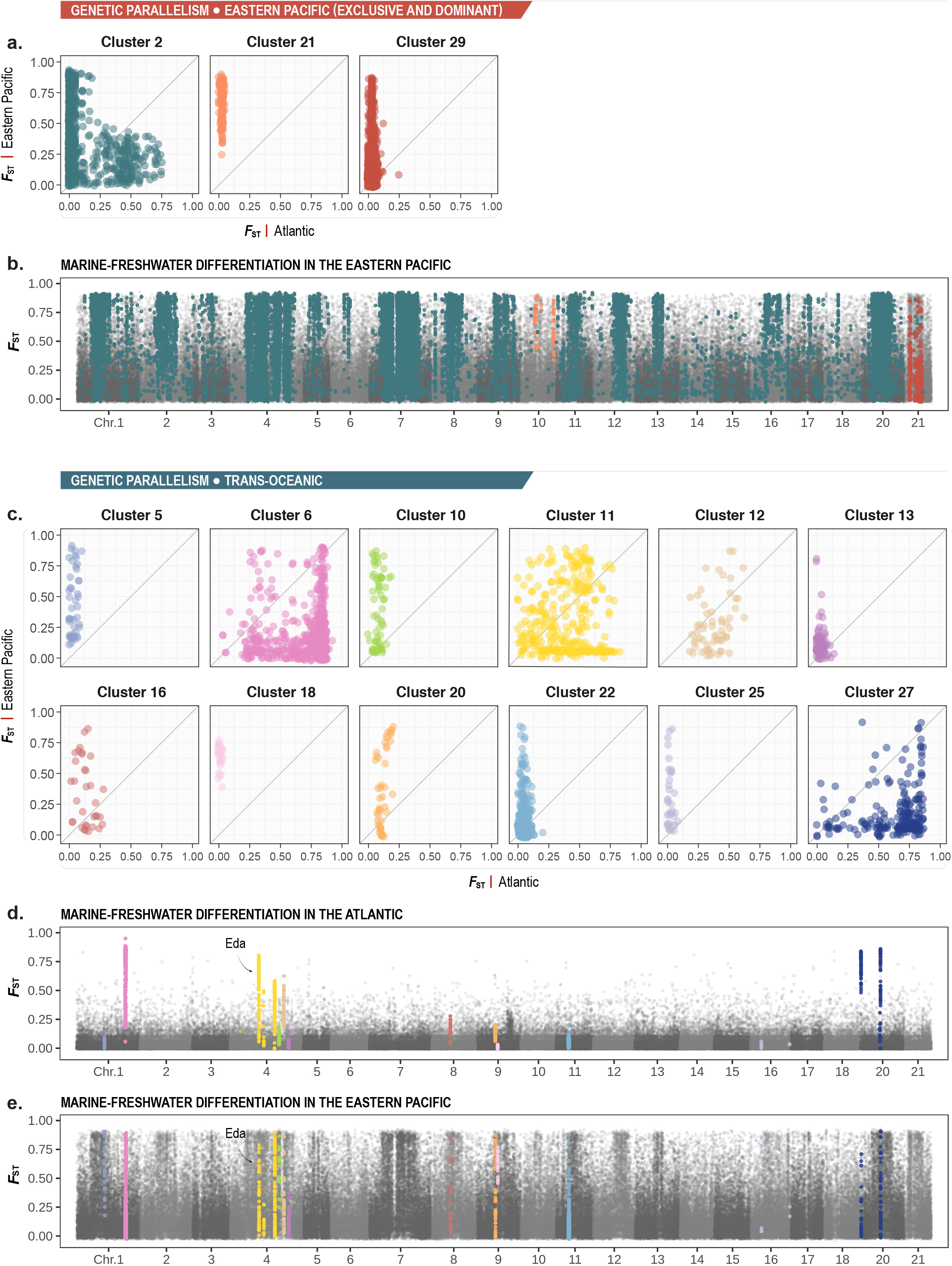
Genetic parallelism identified by the unsupervised and supervised methods. (a) Comparison of marine-freshwater differentiation (*F*_ST_) in the Atlantic (x-axis) and Eastern Pacific (y-axis) datasets for the three LD-clusters (LD-clusters 2, 21 and 29) associated with strong marine-freshwater parallelism in the Eastern Pacific. (b) Genome-wide *F*_ST_ of the Eastern Pacific samples for loci of the LD-clusters coloured as in (a). (c) The same as (a) but for the twelve LD-clusters (5, 6, 10, 11, 12, 13, 16, 18, 20, 22, 25 and 27) that are involved in global marine-freshwater genetic parallelism. (d) and (e) Genome-wide *F*_ST_ of the Atlantic and Eastern Pacific samples, respectively, with colours corresponding to LD-cluster loci in (c). The position of the Ectodysplasin (EDA) locus is indicated in (d) and (e).

**Figure 3.**
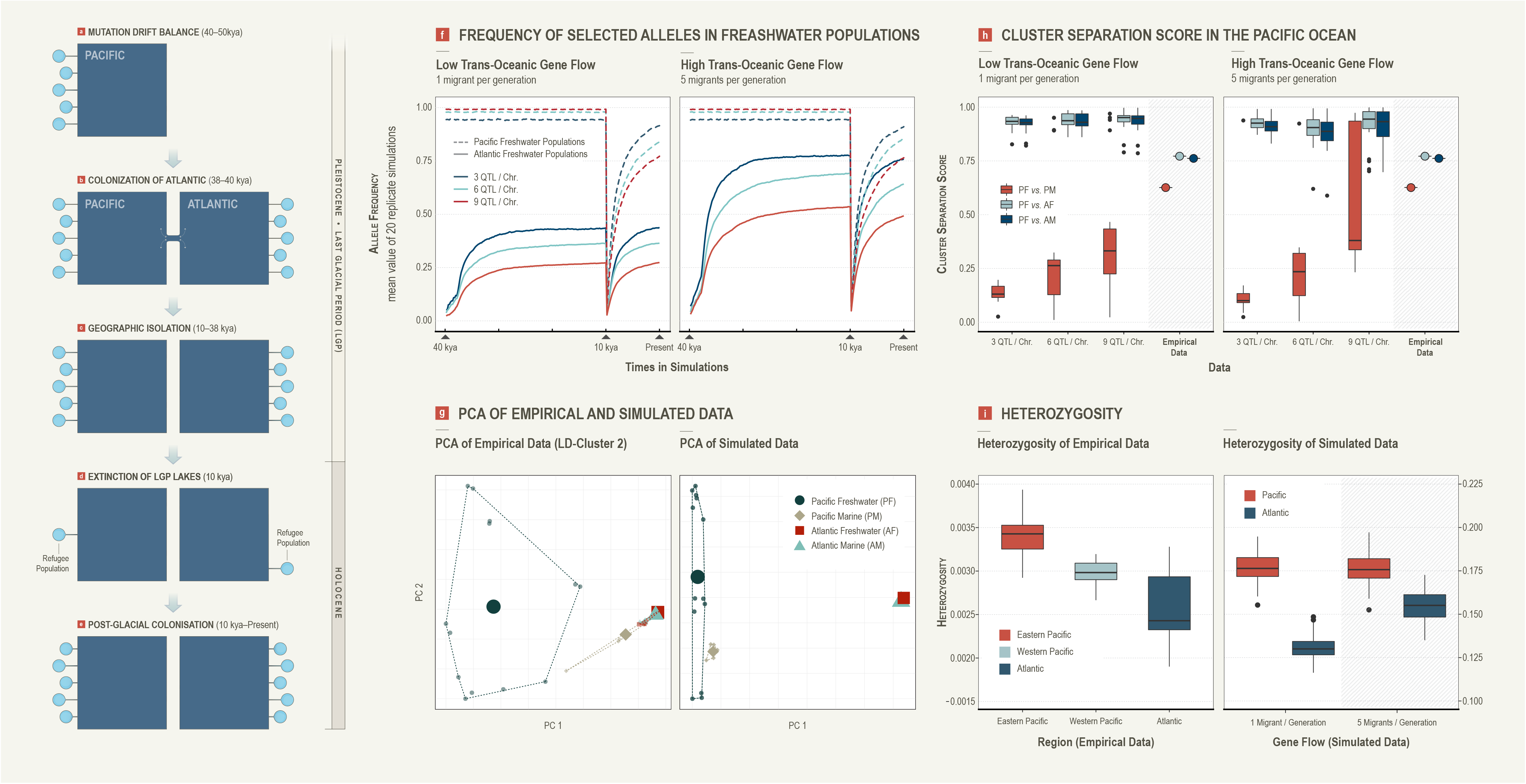
Ecological genetics in simulated data. (a-e) A schematic of the demographic scenario used for the simulations that is consistent with the “transporter hypothesis” of Schluter & Conte (2009). (a) Initial local adaption of the freshwater populations in the Pacific. (b) The colonization of stickleback populations from the Pacific to the Atlantic. (c) Geographic isolation between the two oceans. (d) Extinction of lakes during the last glacial period (LGP) with the survival of refuge populations. (e) The post-glacial colonization of the new freshwater populations. (f) Frequency of selected (freshwater-adapted) alleles in the newly established freshwater populations through generations at high and low levels of trans-oceanic gene flow and different QTL-densities. (g) PCA of the empirical data (LD-cluster 2; left) and the simulated data (right), with ecotypes and geographical regions as shown in the figure legend. (h) Cluster separation score (CSS) of the empirical and simulated data in the Pacific and Atlantic oceans, respectively. (i) Observed heterozygosity in different geographical regions in the empirical and simulated data (Supplementary Information 3).

LDna successfully recovered most of the previously identified regions from Jones et al. (2012) that differentiated marine from freshwater ecotypes, and failed to recover only small regions that had low coverage and relatively low levels of marine-freshwater differentiation (Supplementary Information 5 and Supplementary Fig. 3).

### Trans-oceanic marine-freshwater parallelism

LD-clusters 5, 6, 10, 11, 12, 13, 16, 18, 20, 22, 25 and 27 (a total of 2,502 loci, 0.100% of the dataset; see four representatives in Fig. 1e-h, and Supplementary Fig. 2, Table 3) grouped multiple freshwater individuals from different geographic regions across the Pacific and Atlantic Oceans (for all *P* < 0.05, permutation test for ecotype differentiation), reflecting genetic marine-freshwater parallelism on a global (trans-oceanic) scale. Within those, LD-clusters 6, 11, 12, 16 and 27 (a total of 1,639 loci, 0.065% of the dataset) similarly showed high marine-freshwater differentiation (*F*_ST_, Fig 1e-h, Fig. 2c-e) in both the Eastern Pacific and Atlantic, further suggesting global parallelism. Particularly, loci from LD-cluster 11 mapped to four distinct regions on Chr. V of which one (Fig. 2c-e) mark the position of the *Ectodysplasin* (EDA) gene that is famously responsible for marine-freshwater differences in lateral armour plate development worldwide(Colosimo *et al*. 2005). In contrast, although the remaining clusters (5, 10, 13, 18, 20, 22, 25, a total of 863 loci, 0.034% of the dataset) also grouped freshwater individuals from both the Pacific and Atlantic Oceans (similar to LD-cluster 29 above; Fig. 2c-e and Supplementary Fig. 2), they showed much less marine-freshwater differentiation in the Atlantic Ocean than in the Eastern Pacific (Fig. 2c-e). Among the LD-clusters associated with marine-freshwater differentiation, LD-clusters 6 and 22 covered previously known chromosomal inversions on Chr. I and XI, respectively (Jones et al., 2012, Fig. 1e, Supplementary Fig. 2 and Supplementary Table 3). In addition, we also found a putative novel inversion on Chr. V (LD-cluster 19,241 loci) that was not associated with marine-freshwater differentiation (Supplementary Fig. 2 and Table 3). Neither LDna nor *F*_ST_ analyses could detect any regions of marine-freshwater differentiation in the Western Pacific (Supplementary Fig. 4) and thus, this region is not considered further here.

### Eastern Pacific marine-freshwater parallelism

LD-clusters 2 (53,785 loci, 2.141% of the dataset, Fig. 1b) and 21 (183 loci, 0.007% of the dataset, Fig. 1c) separated Eastern Pacific freshwater individuals exclusively from the remaining samples, a pattern that is not expected by chance alone (permutation test *P* < 0.001, Supplementary Fig. 2, Supplementary Table 3). Rather, this reflects a shared adaptive response among Eastern Pacific freshwater populations. The exception was two freshwater individuals from the Eastern Pacific (ALA; Alaska) that did not group with the other freshwater individuals from the Eastern Pacific but instead with Atlantic (marine and freshwater), Western Pacific (marine and freshwater) and the marine individuals from Eastern Pacific (Fig. 1c, Supplementary Fig. 5). LD-cluster 29 (2,728 loci, 0.109% of the dataset) covering a known inversion on Chr. XXI (Fig. 1d) grouped the Eastern Pacific freshwater individuals (except the two Alaskan individuals above) together with six Atlantic freshwater individuals and one Eastern Pacific marine individual. Because this LD-cluster maps to an inversion, the groups also represent putative inversion karyotypes. Thus, this inversion shows strong ecotype differentiation not only in the Eastern Pacific, but also in a small proportion of individuals outside of the Eastern Pacific that putatively carry the freshwater-adapted karyotype (i.e. the karyotype with the highest frequency among Eastern Pacific freshwater individuals). Notably, no cluster of similar magnitude to LD-cluster 2 – which separates freshwater individuals from one specific region from all remaining samples in the data – could be detected outside of the Eastern Pacific, demonstrating that parallel marine-freshwater differentiation in the Eastern Pacific is much more prevalent than anywhere else in the world.

A small proportion of the loci from LD-cluster 2 (28 SNPs) mapped to regions that showed global parallelism in Jones et al. (2012; Supplementary Table 3). In addition, <1% of all loci in LD-cluster 2 (243 SNPs) showed *F*_ST_ > 0.2 also in the Atlantic (as is evident e.g. from Fig. 2a and Supplementary Fig. 4a). These loci appear to be non-randomly distributed in the genome (Supplementary Fig. 4a), indicating that indeed they are likely to be linked to genomic regions involved in marine-freshwater differentiation in both the Atlantic and the Eastern Pacific. Due to small sample-sizes, the *F*_ST_ Manhattan plots display a considerable amount of noise, particularly in the datasets from the Eastern and Western Pacific Oceans (Fig. 2b,e and Supplementary Fig. 4).

### Geographic structure and regional local adaptation

LD-cluster 1 (10,184 loci, 0.405% of the dataset) separated all Pacific individuals (Eastern and Western) from the Atlantic individuals (Fig. 1a), thus mainly reflecting trans-oceanic geographic structure. LD-clusters 4, 8, 9, 14 and 24 (a total of 526 loci, 0.021% of the dataset, Supplementary Fig. 2) separated freshwater individuals from only one geographic region; this likely reflects geographic clustering but could also contain some loci involved in non-parallel freshwater adaptation. These loci are therefore interpreted as inconclusive with respect to their underlying evolutionary phenomena (Supplementary Table 3). Accordingly, loci from these LD-clusters showed little marine-freshwater differentiation in both the Eastern Pacific and Atlantic (Supplementary Fig. 2), and only 2 loci (from LD-cluster 14) mapped to the *m-f* global regions (Supplementary Table 3). All freshwater individuals in this dataset were important, as they can inform us about the geographic scale of parallel marine-freshwater differentiation (for the LD-clusters where marine-freshwater differentiation also involved geographic regions where marine samples were available).

### Proof of concept simulations

In the simulated data, before *Atl* was colonized from *Pac*, all five freshwater populations in *Pac* were fixed or nearly fixed for the freshwater-adapted alleles of all locally adapted QTL (Fig. 3f). Following the colonization of *Atl* (38-440 Kya; Fig. 3f), the increased frequency of the freshwater allele in the Atlantic freshwater populations depended on both QTL density and the level of gene flow between *Pac* and *Atl* (Fig. 3f). The highest increase in freshwater-adapted alleles in *Atl* was observed when QTL density was low (3 QTL per chromosome) and trans-oceanic gene flow was high (5 migrants/generation, Fig. 3f).

During post-glacial colonization of new freshwater habitats from the sea (10 Kya to present), freshwater-adapted alleles (in both *Pac* and *Atl*) gradually increased in the newly formed freshwater populations (Fig. 3f), reflecting local adaptation. This increase was similarly dependent on the QTL density (both in *Pac* and *Atl*) and trans-oceanic gene flow (only affecting *Atl*, Fig. 3f). These patterns likely reflect the underlying levels of ancestral variation in the sea available for subsequent freshwater adaptation (Supplementary Fig. 6a). The lowest frequencies of freshwater-adapted alleles in the sea were always observed when QTL density was the highest (in both *Pac* and *Atl*) and trans-oceanic gene flow was the lowest (only affecting the *Atl*, Supplementary Fig. 6a). Furthermore, the frequency of freshwater-adapted alleles in both the sea (ancestral variation) and in the post-glacial freshwater populations (local adaptation) depended on whether the QTL were located in low or high recombination regions; the lowest frequencies of freshwater-adapted alleles were always observed in low recombination regions (Supplementary Fig. 6b,c). The freshwater-adapted alleles in both *Pac* and *Atl* freshwater populations never reached similar frequencies during the post-glacial colonization (10 Kya; Fig. 3f) as before post-glacial colonization (>10 Kya; Fig. 3f), showing that ancestral variation in the sea was not sufficient to allow complete local adaptation (i.e. fixation of all original freshwater-adapted alleles) in our simulations. Note that with the rapid fixation of all the freshwater-adapted alleles (that started at frequency 0.5 in the freshwater populations in *Pac*) and the low mutation rate used (1e-8 per site and generation), the contribution of *de novo* mutations (at the QTL) to freshwater adaptation in these simulations are negligible.

In the simulations, present-day marine-freshwater differentiation (mean neutral *F*_ST_) was always low for the two chromosomes without QTL, and high recombination regions of chromosomes that contain QTL (Fig. 4; Supplementary Fig. 6d). In contrast, *F*_ST_ for low recombination regions of QTL-containing chromosomes was high for *Pac* (for all parameter settings), indicating strong islands of parallel marine-freshwater differentiation. This was also true for *Atl* when QTL density was low (3 or 6 QTL per chromosome) and when trans-oceanic gene flow was high, but not when QTL density was high (9 QTL per chromosome; Fig. 4; Supplementary Fig. 6d) and trans-oceanic gene flow was low.

**Figure 4.**
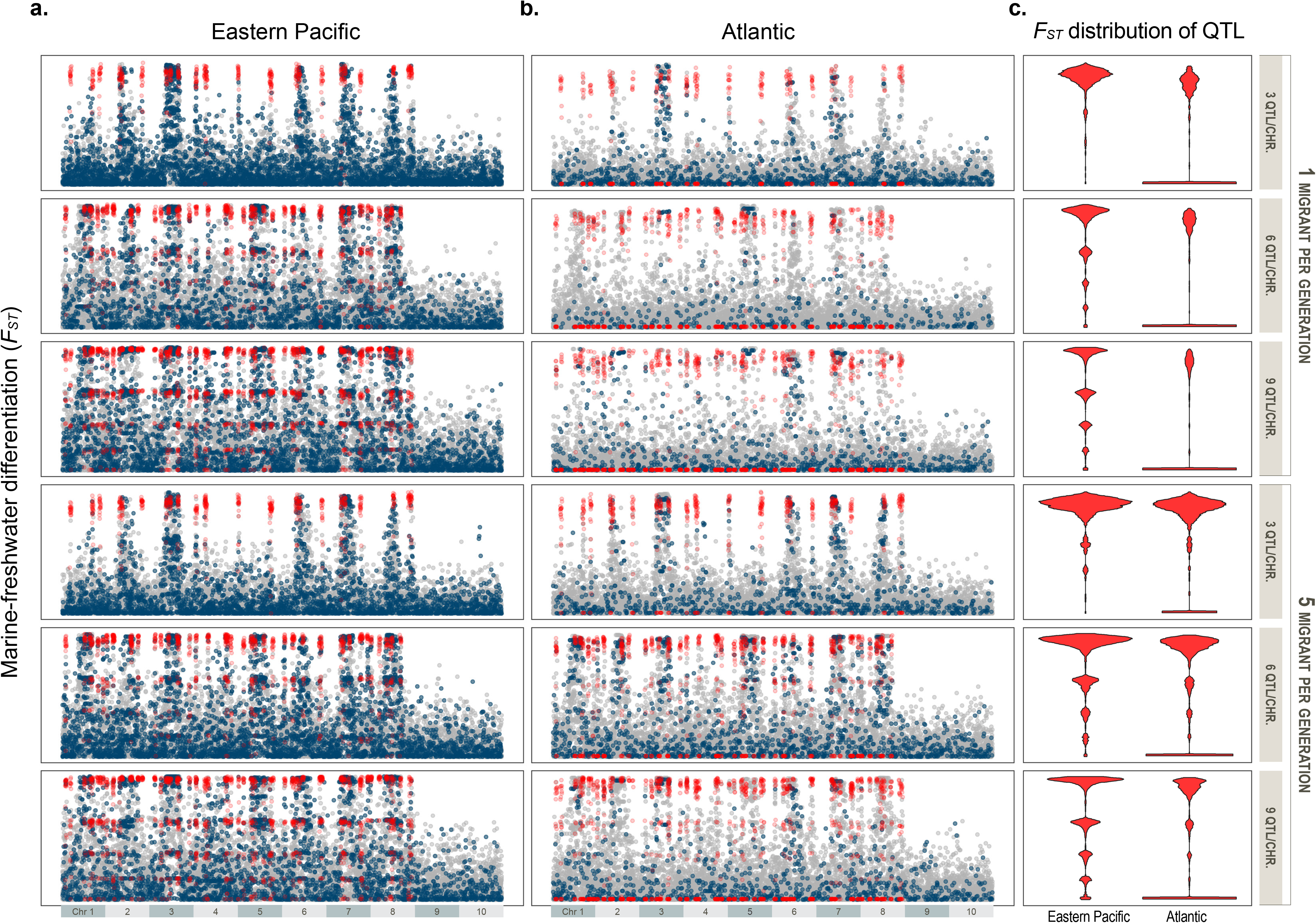
Genomic differentiation in simulated data. (a, b) Genome-wide marine-freshwater differentiation (*F*_ST_) from simulated data (data from the last generation representing present day sampling). For each parameter combination, loci from all 20 replicates were pooled. Red dots indicate QTL, and blue dots indicate loci from LD-clusters that were the most similar to LD-cluster 2 (empirical data) showing the strongest marine-freshwater differentiation in the Eastern Pacific (grey represent non-LD cluster loci). (c) *F*_ST_ distribution of QTL in the simulations (all replicates pooled), indicating the proportion of loci that were either fixed (*F*_ST_~1), lost (*F*_ST_~0), or were fixed to different degrees in only 1, 2 or 3 of the four freshwater populations (0.1 ≲ *F_ST_* ≲ 0.9) since post glacial colonisation. A small amount of noise (along the x-axis) has been added to the QTL positions to improve their visibility.

In the LD-clusters with the strongest *PF* versus non-*PF* differentiation in the simulated data, CSSs were always high (>0.75) between *EF* and the Atlantic populations (*AF* and *AM*), similar to LD-cluster 2 (Fig. 3g-i). However, in contrast to the simulated data, the CSS between *EF* and *PM* for LD-cluster 2 was also high (0.62), whereas for the simulated data this score increased with QTL density (starting from < 0.2), and more so when migration rate during colonization of *Atl* from *Pac* was high (5 migrants/generation; Fig. 3g-i). This is likely due to QTL density increasing the distance between *PF* and *PM* and migration rate decreasing the distance between *Pac* and *Atl* individuals, as CSS here is scaled by the maximum Euclidean distance between any two points in the data. Furthermore, when migration rate was low, no LD-cluster showed any significant CSS between *AF* and *AM*. However, when migration rate was high and with increasing QTL density, LD-clusters similar to LD-cluster 2 were readily observed also in *Atl*. This shows that in simulations, low migration rates and high QTL densities are required to produce patterns similar the observed data.

## Discussion

Using genome-wide SNP data from a comprehensive global sampling of marine and freshwater stickleback ecotypes, we demonstrate that a much smaller proportion of the genome (0.208% of the dataset) is involved in global parallel marine-freshwater differentiation than exclusively in the Eastern Pacific (2.149% of the dataset). This shows that parallel evolution in the three-spined stickleback is much more pervasive in the Eastern Pacific than anywhere else in the world. Indeed, the LD signal from marine-freshwater differentiation in the Eastern Pacific is even stronger than that from geographic structuring between the Pacific and Atlantic Oceans – LD-clusters separating freshwater individuals from the Eastern Pacific comprised five times as many loci than the LD-cluster reflecting geographic structuring between Pacific and Atlantic Oceans. With simulations, we demonstrate that this pattern could partly be explained by the stochastic loss of low frequency freshwater-adapted alleles in the sea during range expansion from the Eastern Pacific. As predicted, the discrepancy between the simulated Pacific and Atlantic populations in both *F*_ST_ and CSS analyses was the highest when trans-oceanic gene flow was low (stronger founder event), but this also required the QTL density of locally adapted loci to be high, as this reduced the levels of standing variation of freshwater-adapted alleles in the Atlantic. However, the loss of ancestral variation due to founder effects and the transporter hypothesis is likely not the only explanation for the large discrepancy in patterns of marine-freshwater differentiation between the Eastern Pacific and Atlantic Oceans. In the following, we discuss alternative biological processes that could potentially contribute to this discrepancy.

### Geographic heterogeneity in standing genetic variation

The “transporter hypothesis”(Schluter & Conte 2009) postulates that a low frequency of freshwater-adapted alleles is maintained in the sea via recurrent gene flow between ancestral marine and previously colonized freshwater populations. This standing genetic variation is what selection acts on during the subsequent colonization of freshwater habitats. This implicitly assumes that a large pool of locally adapted alleles has accumulated over a long period of time, as gene flow is expected to spread potentially beneficial mutations across demographically independent populations (Johannesson *et al*. 2010; Kemppainen *et al*. 2011). In support of this hypothesis, it has been shown that haplotypes repeatedly used in freshwater adaptation are identical by descent (Colosimo *et al*. 2005; Roesti *et al*. 2014) and old – on average, six million years (My), but some are reported to be as old as 15 My (Nelson & Cresko 2018). Notably, these studies analyzed populations from the Eastern Pacific region, which represents the oldest and most ancestral marine population (Fang *et al*. 2020; Fang *et al*. 2018) where three-spined sticklebacks are thought to have persisted since the split from their close relative, the nine-spined stickleback (*Pungitius pungitius*), approximately 26 Mya (Betancur *et al*. 2015; Matschiner *et al*. 2011; Meynard *et al*. 2012; Sanciangco *et al*. 2016; Varadharajan *et al*. 2019). However, populations in the Western Pacific and the Atlantic are much younger, as they were colonized from the Eastern Pacific during the late Pleistocene (36.9-346.5 kya (Fang *et al*. 2019; Fang *et al*. 2020). Furthermore, there is no evidence for trans-oceanic admixture (Fang *et al*. 2019; Fang *et al*. 2020) following the split of Pacific and Atlantic clades, and there are no extant populations of three-spined sticklebacks in arctic Russia between the Kara Sea and the Eastern Siberian Sea. Thus, the spread of freshwater-adapted alleles from the Eastern Pacific to elsewhere via migration through the Bering Strait is unlikely, and has probably not occurred in recent times. Our simulations show that following colonization of freshwater populations from the sea, the accessibility of freshwater-adapted alleles – which is a function of colonization history, QTL-density and recombination rate – largely determines the number of loci that show high marine-freshwater differentiation. Thus, consistent with previous simulations (Feder & Nosil 2010; Roesti *et al*. 2014), genomic islands of differentiation in linked neutral loci require several QTL to cluster in low recombination regions (Fig. 4 and Supplementary Fig. 6d). Furthermore, when trans-Atlantic gene flow was low and QTL density was high, we readily observed LD-clusters that showed high marine-freshwater differentiation only in the Eastern Pacific, not in the Atlantic.

Our simulations and empirical data suggest that both stochastic loss of genetic diversity and selection against freshwater-adapter variants likely played a role in reducing the pool of standing genetic variation of freshwater-adapted alleles in the Atlantic region. During range expansions, genetic diversity is expected to decrease with distance from the source population from which the expansion started (Ramachandran *et al*. 2005). This pattern was very clear in our simulations as well as in the empirical data, both of which show a very clear and statistically significant reduction in heterozygosity in the Atlantic region as compared to the Eastern Pacific region (Fig. 3i, Supplementary Information 3). These results are consistent with some level of founder effect following colonisation of the Atlantic basin from the Eastern Pacific (Supplementary Information 3), which could account for the random loss of standing genetic variation of freshwater adapted alleles Furthermore, since freshwater-adapted alleles are selected against in the sea and thus occur at low frequencies in marine environments, they are even less likely to spread to new geographic regions than neutral alleles (Halliburton & Halliburton 2004; Hyten *et al*. 2006). Consistent with this, the mean individual heterozygosity for LD-cluster 2 loci was 29 times higher in the Pacific compared to the Atlantic Ocean (Supplementary Fig. 7b): a very pronounced difference compared to that in the rest of the genome.

Alternative explanations for the observed discrepancy in patterns of marine-freshwater differentiation between the Eastern Pacific and Atlantic include *i*) stronger spatial genetic structure in marine populations outside of the Eastern Pacific causing heterogeneity in standing genetic variation available for freshwater adaptation, and *ii*) heterogeneity in selective regimes among freshwater habitats, both between Atlantic and Eastern Pacific Oceans and between different geographic areas in the Atlantic. We have further tested these hypotheses but found little or inconclusive support in our data and in other studies (Supplementary Information 3).

### secondary contact in the Eastern Pacific

All of the above hypotheses assume that the original source of the ancestral variation in the Eastern Pacific and elsewhere is the same. That is, ancient Eastern Pacific marine populations carried most ancestral variation of freshwater adapted alleles at low frequencies, sourcing the Atlantic region with freshwater adapted alleles which were partially lost either stochastically or due to selection. However, an alternative hypothesis is that modern Eastern Pacific freshwater variants were not present in the marine ancestors but rather in land-locked, ice-lake freshwater populations (Bierne et al. 2013). As glaciers melted, those populations could have followed meltwater downstream, establishing freshwater populations with a different stock of alleles. Secondary contact with marine sticklebacks during this time might have eroded genetic differentiation across most of the genome, with the exception of those regions involved in freshwater adaptation. In this scenario, the standing genetic variation responsible for Eastern Pacific freshwater adaptation may not have entered the Eastern Pacific marine population until after the end of the last glaciation (i.e. after the closing of the Bearing Strait), with no potential for gene flow to the Atlantic.

There is ample evidence for large ice-lakes during the last glacial period (LGP) in North America, with little (if any) connection to the sea (Baker & Bunker 1985; Bretz 1969; Oviatt 2015; Upham 1896). Thus, a large part of the genetic variation underlying marine and freshwater adaptation in the Eastern Pacific could in principle have evolved in allopatry i.e. separately among the freshwater ice-lake populations and in the sea (and any other potential freshwater water bodies the sea is in contact with). Since the existence of these ice-lakes preceded the invasion of the Atlantic Ocean (Baker & Bunker 1985; Oviatt 2015) the freshwater-adapted alleles potentially residing in such ice-lakes could not have easily spread to the Atlantic. Consistent with this hypothesis is the strong pattern of long-range LD observed among Eastern Pacific marine individuals (Hohenlohe *et al*. 2012), as well as our LDna results which revealed one large cluster separating the Pacific and Atlantic Ocean individuals, and one that specifically separates all Eastern Pacific freshwater individuals from all other individuals. The secondary contact hypothesis is also consistent with close to zero *HE* among Atlantic marine individuals observed for LD-cluster 2 loci (Supplementary Fig. 7a). Curiously, we found significant isolation by distance in the Atlantic but not in the Eastern Pacific where overall population structuring was nevertheless higher than in the Atlantic (Supplementary Fig. 7d). This could be consistent with the secondary contact hypothesis, if introgression was stronger in some regions of the Eastern Pacific compared to others. However, further empirical and simulation studies are needed to test the extent to which this secondary contact hypothesis provides a better explanation for the observed data than the transporter hypothesis alone.

### Conditions that allow global parallelism

Genomic islands of parallel ecotype divergence were more likely to arise in the simulations when several QTL clustered in the same low recombination region. Surprisingly these were also the QTL where the frequency of the freshwater-adapted allele showed the lowest frequencies in the sea and thus, were least likely to spread to Atlantic during colonisation from Pacific. Since QTL in low recombination regions are less likely to be separated by recombination when freshwater-adapted individuals migrate to the sea, it is reasonable to assume that the selection pressure against these “haplotypes’ in the sea is stronger (Hohenlohe et al., 2012). However, this is not consistent with the empirical data showing that the genomic regions most likely to show global parallel ecotype divergence are inversions, where recombination in heterokaryotypes is particularly restricted. Our simulations assume that freshwater-adapted alleles are selected against in the sea (and the strength of this selection is equal for all QTL) while in reality, selection against some of the “freshwater haplotypes/karyotypes” in the sea may be weak or even absent, allowing them to easily spread during range expansions. Consistent with this reasoning, in PCAs based on loci from LD-clusters corresponding to inversions (LD-clusters 6, 22 and 29) several marine individuals also cluster with the freshwater individuals (Fig. 1d,e, Supplementary Fig. 2), indicating frequent occurrence of the “freshwater karyotypes” in the sea. Indeed, Terekhanova et al. (2019) found that the genomic regions most commonly involved in local adaptation in multiple independent freshwater populations were also those with the highest frequencies in the sea. In other words, the most geographically widespread genomic regions involved in freshwater adaptation (*sensu* the transporter hypothesis) are likely to experience the weakest selection against them in the sea, allowing them to remain at higher frequencies in the sea as standing genetic variation (Terekhanova *et al*. 2019).

### Are three-spined sticklebacks a representative model to study parallel evolution?

Since the pattern of parallel genetic differentiation between marine and freshwater stickleback ecotypes in the Eastern Pacific is in stark contrast to what is seen across other parts of the species distribution range, it is reasonable to question the generality of the findings from the Eastern Pacific stickleback studies with respect to parallel evolution on broader geographic scales. A recent review of parallel evolution suggests that even dramatic phenotypic parallelism can be generated by a continuum of parallelism at the genetic level (Bolnick et al., 2018). For instance, the coastal ecotypes of *Senecio lautus* exhibit only partial reuse of particular QTL among replicate populations (Roda et al., 2017), and genetic redundancy frequently underlies polygenic adaptation in *Drosophila* (Barghi et al., 2019). Similarly, using *F*_ST_ outliers to detect putative genomic targets of selection, Kautt et al. (2012, cichlid fishes), Le Moan et al. (2016, anchovy) and Westram et al. (2014, periwinkles) showed that phenotypically very similar populations often share only a small proportion of their *F*_ST_ outliers.

One exception that seems more general across taxa is the repeated involvement of chromosomal inversions in parallel evolution. Chromosomal inversions could store standing variation as a balanced polymorphism and distribute it to fuel parallel adaptation (Morales et al., 2019). For instance, the same Chr. I inversion involved in global marine-freshwater differentiation in three-spined sticklebacks (Jones et al., 2012, Terekhanova et al., 2014, 2019, Liu et al., 2018, this study) also differentiates stream and lake ecotypes in the Lake Constance basin in Central Europe (Roesti *et al*. 2015). Two other clear examples where most genetic differentiation between ecotypes at larger geographic scales is partitioned into inversions come from monkey flowers (*Mimulus guttatus;* Twyford and Friedman, 2015) and marine periwinkles (*Littorina saxatilis;* Faria et al., 2018; Westram et al., 2018).

While our study focuses exclusively on marine-freshwater ecotype pairs of three-spined sticklebacks, other ecotype pairs within freshwater habitats, such as stream *vs*. lake and benthic *vs*. limnetic, also exist. A recent study focusing on stream-lake populations found that putative selected loci showed greater parallelism in the Eastern Pacific (Vancouver Island) than the global scale (North America and Europe; Paccard et al., 2020), i.e. a similar pattern as reported by our study. Furthermore, Conte et al. (2015) studied the extent of QTL reuse in parallel phenotypic divergence of limnetic and benthic three-spined sticklebacks within Paxton and Priest Lakes (British Columbia), and found that although 76% of 42 phenotypic traits diverged in the same direction, only 49% of the underlying QTL evolved in parallel in both lakes. For highly parallel traits in two other pairs of benthic-limnetic sticklebacks, only 32% of the underlying QTL were reported to be shared (Conte *et al*. 2012). Thus, these studies are in stark contrast to the original conclusions of widespread genetic parallelism in three-spined sticklebacks. Notably, the two freshwater individuals from the Eastern Pacific that did not cluster with the remaining freshwater individuals from the Eastern Pacific (and were subsequently removed from the datasets used for *F*_ST_ genome scans) were from Alaska. These two individuals are also phylogenetically distinct from other freshwater individuals from the Eastern Pacific (Fang *et al*. 2018). One explanation for this could be that the highly divergent freshwater populations in the Eastern Pacific have a different colonization history than the Alaskan lakes. More specifically, the former could have been colonized from some divergent ice-lake refugia (see above), whereas the latter could have independently been colonized from the sea.

## Conclusions

Our results demonstrate that genetic parallelism in the marine-freshwater three-spined stickleback model system is in fact not as pervasive as some earlier studies focusing on Eastern Pacific populations have led us to believe. Our analysis of geographically more comprehensive data, with similar and less assumption-burdened methods as used in earlier studies, shows that the extraordinary genetic parallelism observed in the Eastern Pacific Ocean is not detectable elsewhere in the world (e.g. Atlantic Ocean, Western Pacific Ocean). Hence, the focus on the Eastern Pacific has generated a perception bias – the patterns detected there do not actually apply to the rest of the world. Furthermore, our simulations show that the spread of freshwater-adapted alleles can be hampered if colonization of the Atlantic from the Pacific was limited, particularly for QTL clustered in low recombination regions (i.e. those most likely to result in parallel islands of ecotype differentiation). Therefore, geographic differences in the incidence and pervasiveness of parallel evolution in three-spined sticklebacks likely stem from geographic heterogeneity in access to, and amount of, standing genetic variation, which in turn has been influenced by selection as well as historical population demography. Such historical demographic factors include founder events as well as the potential accumulation of genetic ecotypic differences in allopatry during the last glacial maximum, followed by a secondary contact only after the Atlantic Ocean was colonized via the sea from the Eastern Pacific. Hence, while striking genome-wide patterns of genetic parallelism exist (e.g. in Eastern Pacific sticklebacks), the conditions under which such patterns can occur may be far from common, perhaps even exceptional.

## Acknowledgements

We are grateful to the following people who helped in obtaining the samples used in this study: Jacquelin DeFaveri, Anders Adill, Windsor Aguirre, Theo Bakker, Alison Bell, Mike Bell, Bertil Borg, Fredrik Franzén, Akira Goto, Andrew Hendry, Gabor Herczeg, Frank von Hippel, Aki Hirvonen, Jenni Hämäläinen, Markku Kaukoranta, Agnieszka Kijewska, David Kingsley, Yoshinobu Kosaka, Lotta Kvarnemo, Dmitry Lajus, Tuomas Leinonen, Arne Levsen, Scott McCairns, Antoine Millet, Jola Morozinska, Corey Munk, Hannu Mäkinen, Arne Nolte, Kjartan Østbye, Wäinö Pekkola, Jouko Pokela, Mark Ravinet, Katja Räsänen, Dolph Schluter, Mat Seymor, Takahito Shikano, Per Sjöstrand, Garrett Staines, Björn Stelbrink, Ilkka Syvänperä, Anti Vasemägi, Mike Webster, James Willacker, Helmut Winkler, and Linda Zaveik. Our research was supported by grants from Academy of Finland (250435, 263722, 265211 and 1307943 to JM and grant 316294 to PM), the Finnish Cultural Foundation (grant 00190489 to PK) and the Chinese Scholarship Council (grant 201606270188 to BF). We wish to thank three anonymous referees for constructive criticism, and Jacquelin De Faveri for feedback and linguistic corrections.

## Author contributions

PK and JM conceive the concept of the study, with developments from PM and BF. BF, PK and PM performed analyses. PK, BF and PM wrote the manuscript. XF contributed to lifeover analysis. BF visualised the data. JM contributed to shaping the research and manuscript. All authors accepted the final version.

