## Supplementary materials for "Oceans apart: Heterogeneous patterns of parallel evolution in sticklebacks"

### CONTENTS

#### *SUPPLEMENTARY INFORMATION*


**1 |** Three-step linkage disequilibrium network analyses with complexity reduction

**2 |** Testing for LD-cluster association with parallel marine-freshwater divergence

**3 |** Genetic variation and Isolation-by-distance (IBD)

**4 |** Proof of concept using simulated data

**5 |** The power of LDna to detect marine-freshwater differentiated regions


#### *SUPPLEMENTARY FIGURES*


**Supplementary Figure 1 |** Mercator projection of global three-spined stickleback populations used in the study.

**Supplementary Figure 2 |** Visualization of LD-clusters identified by LDna

**Supplementary Figure 3 |** Ability of LDna to recover marine-freshwater differentiated regions from Jones et al. (2012).

**Supplementary Figure 4 |** Genome-wide marine-freshwater differentiation ($F_{ST}$) in the Atlantic, Eastern Pacific and Western Pacific Oceans.

**Supplementary Figure 5 |** PCA plot of LDna clusters with population identification.

**Supplementary Figure 6 |** Allele frequency changes in simulated data.

**Supplementary Figure 7 |** Population variation and Isolation-by-Distance (IBD) in marine three-spined stickleback populations.

**Supplementary Figure 8 |** Network tree of the 29 clusters identified by LDna.

**Supplementary Figure 9 |** Recombination map of simulated chromosomes.

#### *SUPPLEMENTARY TABLES*

**Supplementary Table 1 |** Detailed sample information.

**Supplementary Table 2 |** Specification of different sampling schemes for the $F_{ST}$ analyses between marine and freshwater populations.

**Supplementary Table 3 |** Summary of all LD-clusters.

### SUPPLEMENTARY INFORMATION

#### INFORMATION 1 | Complexity reduction using three-step linkage disequilibrium network analysis

Linkage disequilibrium network analysis (LDna) is an efficient way to partition loci into sets that represent the same phylogenetic pattern in population genomic data[1,2]. Since it relies on LD estimates for all pairwise comparisons between loci in the dataset, LDna is currently limited to ~20 000 SNPs at a time. However, with high-density SNP data, many adjacent loci are likely not independent and display high LD due to physical linkage. It is thus reasonable to initiate LDna analyses by first grouping loci within small windows within each chromosome, and letting only one locus represent each cluster[2]. This approach has previously been used to successfully reduce complexity of population genomic data to increase the power of genome-wide association analyses[2], and is now part of the R Package LDna[1]. To produce the input for such LD-network clustering, we estimated pairwise LD between loci from genotype likelihoods using the program ngsLD[3]**,** setting the minimum SNP minor allele frequency to 0.05. To this end, the original "LDnClustering" function was modified such that it now allows LD values to be estimated by other software and subsequently imported as edge lists (function "LDnClustering_EL"; LDna v.0.65). For this initial step we only considered pairwise LD values within the closest 100 SNPs (w2=100) along each chromosome, and performed LD clustering within non-overlapping windows of ~1,000 SNPs (w1=1000). This size naturally varies slightly, as window break points were only placed in regions where LD between adjacent loci were below the threshold of 0.5 (thershold1=0.5) for at least 10 consecutive SNPs along the chromosomes (w1=10). Previous unsupervised methods such as Hidden Markov Model (SOM/HMM) in Jones et al.[4] find genomic regions that support a certain phylogenetic tree, thus each of these genomic regions are likely to harbour at least some SNPs in high LD that are diagnostic for each tree. Because of this, and to reduce the number of pairwise comparisons in our analyses, we excluded "singleton SNPs"[2] defined as loci with the maximum pairwise LD value within the 1,000 SNP windows < 0.8 (threshold2=0.8). From each cluster (containing at least two loci), we chose the SNP exhibiting the highest median LD with all other SNPs in the same cluster ($LD_{MED}$), which we refer to as Maximally

Connected Loci (MCL), to represent the whole cluster. When only two loci were available, one was randomly chosen as the MCL.

In the second step, LDna analyses were performed for each chromosome separately using the MCL. LDna starts by producing a single linkage clustering (SLC) tree (a hierarchical clustering algorithm which combines two clusters connected to each other by at least one edge) based on a pairwise matrix of LD (in this case, LD values between MCL). As the LD threshold is sequentially lowered, an increasing number of loci will be connected to each other in a fashion that reflects the similarity of their phylogenetic signals. For each cluster merger (with decreasing LD threshold) the change in median LD between all pairwise loci in a cluster before and after the merger is estimated as λ. When two highly interconnected clusters merge, λ will be large, unlike when only a single locus is added to an existing cluster. The large λ signifies that these two clusters bear distinct phylogenetic signals. Here we set the parameter $\lambda lim$=2 to extract only clusters with λ above 2, and set the parameter $|E|min$ = 30, which requires a cluster to have at least 30 edges (also limiting the minimum size of the cluster to seven SNPs). Thus, clusters with high λ (relative to other clusters in the clustering tree) and many interconnected loci signify "outlier clusters". While the above settings are arbitrary, generally a wide range of parameter values result in the same clusters being extracted when the LD signal is strong[1]. Because of this, we also considered all "compound outlier clusters" (COCs), i.e. those that contain clusters further up the single linkage clustering tree that have already been identified as "single outlier clusters" (SOCs). We then manually identified outlier clusters that signified major mergers in the single linkage clustering tree (joining of two distinct branches), as demonstrated in Supplementary Fig. 8. This was necessary since no single combination of $|E|min$ or *phi* could identify these mergers consistently within the single linkage-clustering tree, either for each chromosome separately or in the single linkage-clustering tree among the trees from the different chromosomes (see tutorial provided in the LDna package on guidelines for further details). This maximized the number of loci that represented distinct phylogenetic signals in the datasets, while avoiding over-merging of clusters that actually contain two or more distinct genetic patterns (see advanced LDna tutorial for details: https://github.com/petrikemppainen/LDna/blob/master/vignettes/LDna_advanced.pdf).

In the third step, we extracted a single MCL from each of the clusters identified in the previous step, and used these loci to group outlier clusters from the different chromosomes (by LD) into global LD clusters (Supplementary Fig. 8). From the resulting clusters, we then traversed hierarchically back through the clusters produced in this three-step LDna approach to the original loci.

**INFORMATION 2 | Testing for LD-cluster association with parallel marine-freshwater differentiation**

Linkage disequilibrium network analyses (LDna) finds clusters of highly correlated loci in population genomic datasets. These correlations, as estimated by linkage disequilibrium (LD, see Supplementary information 1), can be caused both by non-neutral and neutral processes, most notably population structuring (neutral), inversions (neutral or non-neutral) and parallel ecotype divergence (non-neutral). If the loci from an LD-cluster that is formed by parallel marine-freshwater differentiation are used in a PCA, the expected result is that freshwater individuals will be separated from all other individuals in that geographic region. Thus, to determine if groupings of individuals in a PCA based on loci from a given LD-cluster is reflecting parallel marine-freshwater differentiation, we first need to identify sets of individuals in a PCA that contains a higher proportion of freshwater individuals that expected by chance. As some geographic locations lack marine individuals in our dataset, such groups of freshwater individuals (here referred to as the ’in-group’ individuals), if caused by geographic structuring, by necessity will separate all freshwater individuals from that location from all other locations. Thus, to determine if groupings of individuals in a PCA based on loci from a given LD-cluster is reflecting parallel marine-freshwater differentiation, we first need to identify sets of individuals in a PCA that contain a higher proportion of freshwater individuals that expected by chance. This is because the probability that the in-group only contains freshwater individuals is high even if allelic correlations in the LD-cluster is caused by geographic structuring (and not freshwater adaptation), if no marine individuals from that location exist in the data. These two steps are detailed below.

*Identifying in-groups*

It is quite possible to visually identify sets of individuals within a PCA that contain a high proportion of freshwater individuals, if such groups are clear. However, in an attempt to be as objective and consistent as possible across all LD-clusters, we followed the protocol specified below (custom R-script available from Dryad) to determine potential in-groups. First, we used a model-based clustering algorithm in which the models are estimated by an Expectation Maximization (EM) clustering algorithm. This algorithm is initialized by hierarchical model-based agglomerative clustering (function 'MClust', from R-package 'mclust'[5]), where the optimal model is selected according to the Bayesian Information Criterion (BIC) using uniform priors and default settings. From the optimal clustering solution (containing a minimum of two clusters), we then divided the individuals into two clusters (if the initial clustering solution contained more than two clusters) based on hierarchical clustering (function 'hclust', R-package **'**ape'[6], using the 'median' method) on the Centroids of the initial clusters. This ensured that a potential in-group always contained individuals that were the most 'closely related'. We then only considered clusters with a minimum of seven individuals and 85% freshwater individuals as potential in-groups. Finally, we kept the clusters that deviated the most from the numbers of freshwater and marine individuals expected by chance (as estimated by $X^2$). We further included freshwater individuals closest to the convex hull enclosing the primary in-group if 1) the convex hull after including this individual did not also include marine individuals and 2) if the freshwater individual was no further from the centroid of the primary in-group than three times that of the closest (to the same centroid) marine individual (as otherwise it should not belong to the in-group). We repeated this 'hull expansion' procedure with the second closest freshwater individuals until the conditions above were no longer met, producing a secondary in-group. Note that hull expansion did not always occur. As a last step, if the final convex hull surrounding the primary in-group contained any marine individuals, these were removed as long as its distance to the in-group centroid was no longer than three times the distance from the secondary in-group centroid to the closest freshwater individual outside of the hull (as then, it clearly should be included in the in-group). If the model-based clustering did not produce any in-groups, we initiated the procedure with hierarchical clustering as above instead of the model-based clustering, which was only necessary for LD-cluster 29. All LD-clusters for which in-groups were not found were not considered further as potentially being caused by parallel marine-freshwater differentiation.

*Testing for marine-freshwater association of in-groups*

We then used a permutation approach to test whether the in-group contained more freshwater individuals than expected by chance, taking into account geographic sampling. First, all individuals were separated into seven geographic groups as specified in main text (Eastern Pacific [EP], Western Pacific [WP], Western Atlantic [WA], White and Barents Seas [WB], British Isles and North Sea [UK], Baltic Sea [Bal] and Norwegian Sea [Nor][7]; Fig. 2j). Two of these were from the Pacific Ocean and the remaining five from the Atlantic. An $X^2$ model fit was estimated by a log-linear model using the function 'loglm' from R-package 'MASS', with formulae specified as "*~Grp+Ecotype+Location*" where Grp[in, out] refers to whether the individuals belong to an in-group or not, Ecotype[marine, freshwater] refers to whether an individual is a marine or freshwater ecotype, and Location[WP, EP, Bal, Nor, UK, WB, WA], and refers to the individuals location. Since this is not a valid test *per se*, we used the $X^2$ estimate as a test statistic and generated a null-distribution by permuting the in-group assignment in the dataset within each location (n=$10e^5$) to obtain a *P*-value.

**INFORMATION 3 | Genetic variation and Isolation-by-distance (IBD)**

Under the hypothesis that the colonization of the Atlantic from the Pacific involved a limited number of founder individuals, genetic diversity (individual heterozygosity, H) is expected to be lower in the Atlantic compared to the Eastern Pacific[8]. To evaluate this hypothesis, we classified marine individuals in our data into three geographic regions: Eastern Pacific (EP), Western Pacific (WP) and Atlantic (ATL). Since we only had 4 marine individuals with sufficient coverage from the EP, we incorporated 30 randomly selected RAD-sequenced individuals from three Oregon marine populations (n=10 in each) from a published study[9]. Note that since the restriction enzyme used to generate that dataset (SbfI) was not the same as in our study (PstI), they could not be used for our main analyses. Only marine individuals were used for these analyses since the inclusion of freshwater individuals would inflate genetic diversity, especially in the Eastern Pacific.

To calculate *H* we first mapped the raw sequences of EP individuals using the same pipelines as employed for the rest of the data, and then calculated H based on genotype likelihoods across genomic regions with ANGSD. From BAM files, we first estimated the allele frequency likelihood (using the filters -GL 1, -doCounts 1, -setMinDepth 2, -uniqueOnly 1, -remove_bads 1, -minMapQ 20, -minQ 20, -C 50), and then the one-sample site frequency spectrum. Note that the samples from Jones et al. (2012) were excluded in the $H_E$ calculation due to low read depth (≤1). The final H in each sample was calculated as the number of heterozygous sites divided by the sum of monomorphic and heterozygous sites within each individual. We then estimated H of LD-cluster 2 (the cluster showing signature of parallel evolution in EP freshwater populations) across regions. The data from Catchen et al (2013) were produced with the enzyme SfbI, an 8 bp cutter with a recognition site (CCTGCAGG) whose core 6 bp are the recognition site of PstI (CTGCAG). Thus, a RAD library produced with SbfI is expected to sample, on average, 1/16$^{th}$ of the genomic regions sampled in our study. We therefore selected loci within LD cluster 2 that had less than 20% missing data in each region (including EP data from Catchen et al., 2013), after applying the same filters as above, yielding total of 1217 loci. Then we calculated individual heterozygosity from one-sample SFSs as outlined above. The results showed a significant decrease of genome-wide H across the region EP, WP to ATL (GLM, $F_{2,64}$=43.05, $P$<0.001; Fig. 3i, Supplementary Fig. 7a), while heterozygosity of LD-cluster 2 for the ATL populations was close to zero and 29 times lower than in the EP (GLM, $F_{2,64}$=91.9, $P$<0.001; Supplementary Fig. 7b).

We then looked for correlations between pairwise genetic (***F*~ST~**[10]) and geographic (km) distances (isolation-by-distance; IBD). For geographic distance we used the pairwise least-cost distances between marine samples as calculated by the R-package *Marmap*[11]. In the IBD analyses, individuals from the same locality were treated as a population, and only the populations with at least more than two individuals were considered[12]. The Eastern Pacific and Western Atlantic samples in our dataset were not included in IBD analyses due to the low numbers of populations and samples. However, we incorporated $F_{ST}$ and results of EP populations (224 individuals of eight populations) from Morris et al. (2018). Note this RAD dataset was also not included to the main analyses due to different enzyme used. Significance of IBD was tested with a Mantel test permuted 9999 times using the function *mantel* from the R package 'vegan'[13]. IBD was significant in the Atlantic but not in the Eastern Pacific, although population structuring in general was still higher in the Eastern Pacific (Supplementary Fig. 7d).

We have discussed how the above results support the founder effect hypothesis tested by our simulations and the secondary contact hypothesis in the main document. Here we focus on additional alternative hypotheses. For instance, the lack of parallel islands of ecotype differentiation in the Atlantic could be due to stronger spatial genetic structure in marine populations outside of the Eastern Pacific. Indeed, high levels of genetic differentiation among Atlantic marine populations have been reported[14,15]. This structuring would cause heterogeneity in the standing genetic variation available for freshwater adaptation, which was not tested in our simulations. Limited gene flow among marine populations may increase the chance that freshwater-adapted alleles are stochastically lost in some of the sub-populations, thus resulting in smaller and more heterogeneous pools of freshwater-adapted alleles. While there is indeed a significant IBD signal in our Atlantic marine populations (but not in the Eastern Pacific; Supplementary Fig. 7d), spatial structure in the Eastern Pacific seems to be even stronger (Supplementary Fig. 7; data from[16]). Thus, we find no support for the hypothesis that differences in the degree of spatial genetic structuring among marine populations within regions would explain the observed patterns.

Based on results of this and earlier studies, there is also evidence for parallel evolution in Europe[17-20], though such patterns are locally restricted and involve fewer genomic regions than in the Eastern Pacific. Ferchaudand Hansen (2016) and Liu et al.[18] reported little parallelism in genomic regions involved in local adaptation, most notably in Denmark and Greenland, compared to the Eastern Pacific. Similarly, Terekhanova et al.[20] identified a total of 19 distinct genomic regions showing consistent marine-freshwater divergence in the White Sea (later refined to 21 regions[19]), all of which were also identified as differentiation islands between marine and freshwater ecotypes in the Eastern Pacific[4,21]. Thus, while local adaptation in all geographic regions likely stems from standing genetic variation originating from the Eastern Pacific (*sensu* the transporter hypothesis), only Eastern Pacific populations show extensive parallel evolution over a larger geographic area. These results are consistent with our LDna analyses, where no LD-cluster showed parallel marine-freshwater differentiation exclusively among non-Eastern Pacific samples.

Another possible explanation for the lack of globally shared genetic parallelism is heterogeneity in selective regimes among freshwater habitats, both between Atlantic and Eastern Pacific Oceans and between different geographic areas in the Atlantic[22-24].

However, there is currently no data to assess the extent to which differences in selection regimes could contribute to the observed heterogeneity in parallel patterns of marine-freshwater differentiation at the trans-oceanic scale. A recent simulation study showed that the probability of genetic parallelism from standing genetic variation rapidly declines as selection starts to change from fully parallel (optima angle of 0°) to divergent (optima angle of 180°), especially when a large number of traits affect fitness[25]. For example, populations adapting to optima separated by an angle of just 33° might have only 50% of shared beneficial alleles, even if they have access to the same pool of genetic variation. However, in this model, adaptation proceeded via the sorting of naive alleles and not via alleles that are "pretested" by selection (as under the transporter hypothesis) when parallel evolution is particularly likely[25].

#### INFORMATION 4 | Proof of concept using simulated data

We performed simulations using the program quantiNemo[26] based on the following demographic model. Our model is initiated by simulating a single marine population that is connected to five independent freshwater populations by symmetrical gene flow of one migrant per generation, for 20ky (10k generations). This represents the ancestral Eastern Pacific Ocean (*Pac*). A new ocean representing the Atlantic (*Atl*) is then allowed to be colonized from *Pac* between 38 and 40 Kya, with no further trans-oceanic gene flow possible beyond 38 Kya. *Atl* is also connected to five independent freshwater populations. While the freshwater populations in *Pac* are already populated from the simulation start, the freshwater populations in *Atl* can naturally only be colonized following range expansion from *Pac*. To simulate the retreat of the Pleistocene continental ice sheets and the colonization of newly formed freshwater habitats, we firstly removed four of the freshwater populations (at 10 Kya) in both oceans. Immediately following this, four new freshwater locations were allowed to be colonized from the sea while keeping one of the original freshwater populations as a "glacial refugium" that could continue to feed freshwater-adapted alleles to the sea as standing genetic variation. While this process in reality was most likely gradual and not instant as in our simulations, the end result is the same; most Pleistocene freshwater populations are no longer present today (with the exception of potential glacial refugia), and most present day populations were colonized from the sea following the retreat of the Pleistocene ice sheets. Thus, according to the transporter hypothesis, post glacial local adaptation is only possible due to the spread of freshwater-adapted alleles from the sea[27] (Fig. 3a-e, but see Bierne et al.[28] for an alternative scenario). The carrying capacity (K) that equates to $N_e$ under mutation/drift equilibrium was kept at 10,000 individuals in the sea and 1,000 individuals in the freshwater populations. This historical global population demographic scenario and the population sizes are in line with findings from Liu et al.[29] (2016) and Fang et al.[7,30]. Generation time was assumed to be two years. We allowed two different levels of trans-oceanic gene flow during the colonization of the Atlantic – one or five migrants per generation.

From previous studies[31,32], we know that recombination rate variation plays an important role in the formation of differentiation islands around locally adaptive loci. Thus, we configured a genetic map where the recombination distance (as measured in centiMorgans, cM[33]) was shorter at the chromosome centres to simulate the presence of centromeres. Recombination distance versus physical distance between loci can be seen in Supplementary Fig. 9. We simulated ten equally sized chromosomes of a total length of 100 cM. This relatively short total map length was chosen such that fewer neutral markers (n=1,000) would be needed to detect differentiation islands, in the interest of computational speed. To demonstrate the effect of the number of differentially selected freshwater-adapted traits, each coded by a single quantitative trait locus (QTL), we simulated datasets with 24, 48 and 72 QTL equally distributed among the first eight chromosomes (3, 6 or 9 QTL per chromosome). The remaining two chromosomes were left without any QTL. The positions of these QTL within chromosomes were randomly selected (Fig. 4), and then fixed for all simulation replicates (n=20). We also ensured that there was always some recombination distance between a QTL and the nearest neutral locus, although sometimes by necessity this was very small, for example around the simulated centromeres and telomeres. In other words, no QTL position was exactly the same as any of the positions for the neutral markers. The allelic effects of the QTL were either zero (allele 1) or 10 (allele 2), with the selection optima in the marine habitat being zero and the selection optima in all freshwater populations being 20. Thus, a freshwater individual homozygous for allele 2 for a given QTL meant that the individual was at its optimal phenotype, and *vice versa* for marine individuals. The selection intensity was set to 100 for the freshwater habitat and 200 for the marine habitat, with lower values translating to stronger selection intensity (see quantiNemo manual for details). Lower selection intensity in the sea allowed higher frequencies of the freshwater allele (allele 2) in the sea as standing genetic variation to facilitate rapid local adaption in the newly formed freshwater populations. Per-site germ-line mutation rate was set to $1.0e^{-8}$ per generation for all loci. Full simulation details can be obtained from Dryad. Population genomic datasets for both neutral loci and QTL were saved every 50 generations. All allele frequencies started at 0.5, thus also initially allowing all QTL to be fixed in the Pacific freshwater populations. We allowed 10,000 generations for *Pac* to reach mutation-drift-selection equilibrium. In the absence of selection, the time to reach mutation/drift equilibrium is $4N_e u$ (where $u$ is the mutation rate). However, for all neutral loci that are linked to selected loci, this process is likely to be much faster, hence the relatively short burn-in time relative to *K*.

The combination of two trans-oceanic gene flow rates (one or five migrants/generation) and three different QTL settings (ca. 3, 6 or 9 QTL per chromosome) resulted in six different parameter settings. Custom R scripts were used to retrieve and parse outputs from the simulations. We monitored the allele frequency of freshwater-adapted alleles for the QTL to assess levels of standing genetic variation at 50-generation intervals in both marine and freshwater habitats. Genome differentiation was estimated for the last generation in the simulations. In each replicate simulation, the four freshwater populations that were colonized after the Pleistocene glaciations were pooled and $F_{ST}$ was estimated for neutral loci between the marine and pooled freshwater populations; this was performed separately for each ocean. The $F_{ST}$ calculations were based only on the neutral loci, thus assuming any genetic signal of selection is due to LD with a QTL.

#### INFORMATION 5 | The power of LDna to detect marine-freshwater differentiated regions

From whole genome sequences of 21 individuals, Jones et al.[4] identified 812 regions showing parallel marine-freshwater differentiation in the Eastern Pacific (tree "i" of Jones *et al.*[4]; hereafter "*i*-regions") and 81 regions showing global parallelism (hereafter "*m-f*" regions). They used a self-organizing map-based iterative Hidden Markov Model (SOM/HMM) to detect genomic islands of marine-freshwater differentiation. This finds regions of the genome that support a given set of commonly found tree topologies in the data, where the loci may or may not be in high LD with each other. Conversely, LDna exclusively depends on LD between pairs of loci. Furthermore, in the first step of LDna (only keeping loci with $r^2>0.8$ within windows of 100 consecutive SNPs), the original 2.5 million unfiltered loci were reduced by 92% to 214k SNPs. The question is: to what extent did this affect the power of LDna to detect genomic regions of marine-freshwater differentiation?

To benchmark our LDna analyses, we quantified the proportion of the marine-freshwater differentiated regions from Jones et al.[4] to which at least one SNP in our datasets mapped. To do so, the coordinates of the Jones et al.[4] genomic regions were converted to the current version of the three-spined stickleback reference genome with CrossMap v0.2.8[34] using the 'liftOver chain' file generated by the program flo (https://github.com/wurmlab/flo) with default settings. SNPs from the original 2.5 million SNPs used for the LDna analyses mapped to 78% and 93% of the *m-f* and the *i*-regions, respectively (Supplementary Fig. 3a). For the LD-cluster loci (71k SNPs) the corresponding numbers were 58% and 71%, respectively. Therefore, in those regions for which we had coverage, 19.75% and 22.72% of the *m-f* and the *i*-regions, respectively, were not successfully recovered by LDna. However, these regions were on average smaller, had lower coverage (less sampled SNPs) and $F_{ST}$ values than the correctly recovered regions (Supplementary Fig. 3b-d). For instance, when taking into account all original 2.5M SNPs, the median number of SNPs in the *m-f* regions was 2.5 times larger than the median number of SNPs in the regions that were not recovered by LDna (4.3 times larger for *i*-regions; Supplementary Fig. 3b). When only considering SNPs that passed the first LDna filtering step (214k SNPs), the corresponding numbers for the *m-f* and *i*-regions were 2.1 and 3 times, respectively. The median size of the *m-f* regions that were correctly recovered by LDna was 2.4 times larger than the median size for regions that were not recovered by LDna (3.8 times larger for *m-f* *i*-regions; Supplementary Fig. 3c). Furthermore, for *m-f* regions, the median $F_{ST}$ (Atlantic dataset) for SNPs in regions that were recovered by LDna was 4.3 times larger than in regions that were not recovered by LDna (1.7 times for the Eastern Pacific data set; Supplementary Fig. 3c). For *i*-regions, $F_{ST}$'s were roughly equal in both the Atlantic and the Eastern Pacific data sets. Thus, when we had low coverage (few SNPs in a region) LDna was not able to identify regions showing weaker signatures of selection.

Among the 52,790 SNPs from LD-cluster 2, 8482 SNPs mapped to the 66% of the *i*-regions from Jones et al.[4]. Thus, while LD-cluster 2 and the *i*-regions undoubtedly reflect the same evolutionary phenomena (marine-freshwater differentiation in the Eastern Pacific), 84% of LD-cluster 2 loci nevertheless did not map to *i*-regions. This is likely because LDna can account for the mosaic-like pattern of LD in population genomic data by grouping loci (within windows) based on LD regardless of what overall tree topology that genomic region reflects (See Fig. 1 in Li et al.[2]). LDna is thus expected to have higher resolution on smaller genomic scales compared to SOM/HMM.

While LDna certainly may have missed small sets of outlier loci or single-SNP outliers, low signal to noise ratio (small sample sizes) and background differentiation means that there are no alternative SNP-based outlier analyses available that could differentiate a true single-SNP $F_{ST}$ outlier (i.e. a unique outlier SNP that is not in LD with many adjacent SNPs) from noise, especially in the Western and Eastern Pacific datasets. However, background differentiation and noise are not expected to lead to high LD among large sets of loci, so LDna is inherently resistant to potential false positives that arise from noise, as it requires a certain number of loci that are sufficiently differentiated from other such groups as measured by $\lambda$ to detect potential outliers. This is why LDna could be conducted on all (2.5 million) unfiltered SNPs (LD estimated from genotype likelihoods), whereas some filtering of the $F_{ST}$ datasets was necessary to reduce background noise. Indeed, a testament to this is that LDna could identify LD-cluster 2 that is unique to the Eastern Pacific, where noise and background differentiation was the highest (due to small sample sizes as well as low sequencing quality).

**Supplementary Figure 1 | Mercator projection of global three-spined stickleback populations used in the study.** 166 three-spined stickleback individuals from 63 localities were used, including 119 freshwater individuals and 47 marine individuals. For a complete list of samples, see Supplementary Table 1.

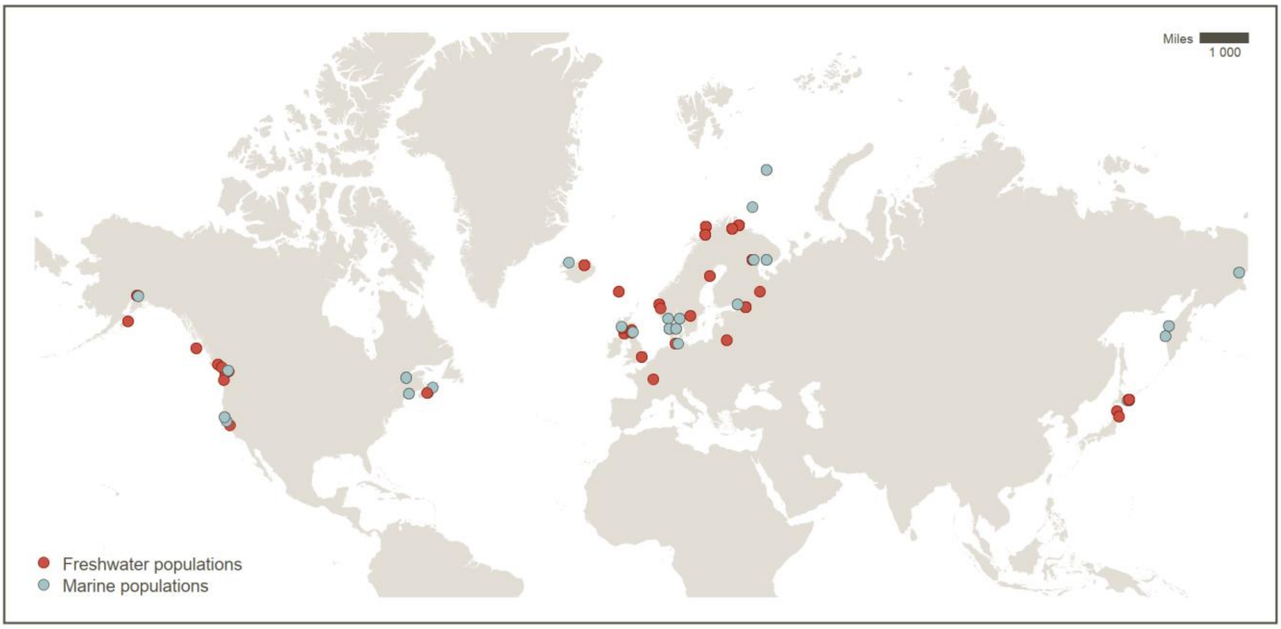

**Supplementary Figure 2 | Visualization of all LD-clusters identified by LDna.** In each panel, I) the top and II) middle plots represent the marine-freshwater differentiation ($F_{ST}$) of the clustered loci of the individuals in the Atlantic and Eastern Pacific, respectively. III) The bottom left plot shows population differentiation based on loci in each LD-cluster (principal component analysis; PCA). Only one chromosome is presented on the x-axis when the clustered loci were located on a single chromosome. IV) The bottom right plot depicts the number of in-group samples (as positive value) and non-group samples (as negative value). Global samples from various regions are shown in different colours; freshwater ecotypes are indicated by light-colour and marine ecotype by dark-colour. The same colour scheme was used in the PCA. The p-values were obtained from permutation tests of cluster separation (Supplementary Information 1).

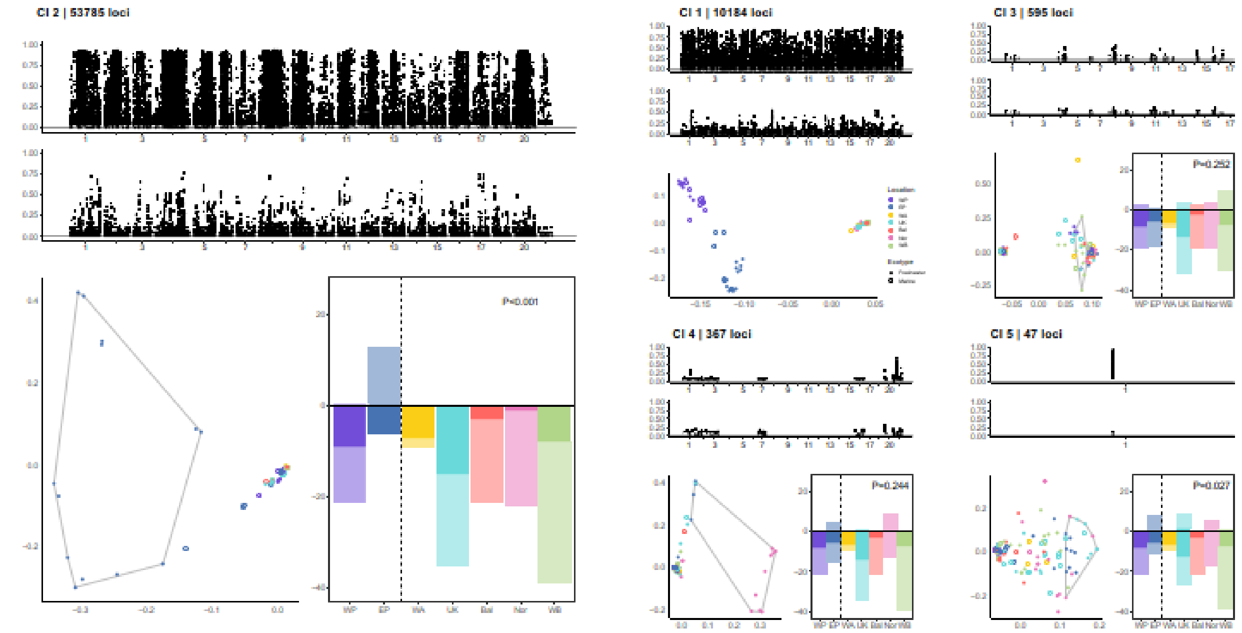
CI 2 | 53785 loci
CI 1 | 10184 loci
CI 3 | 595 loci
CI 4 | 367 loci
CI 5 | 47 loci

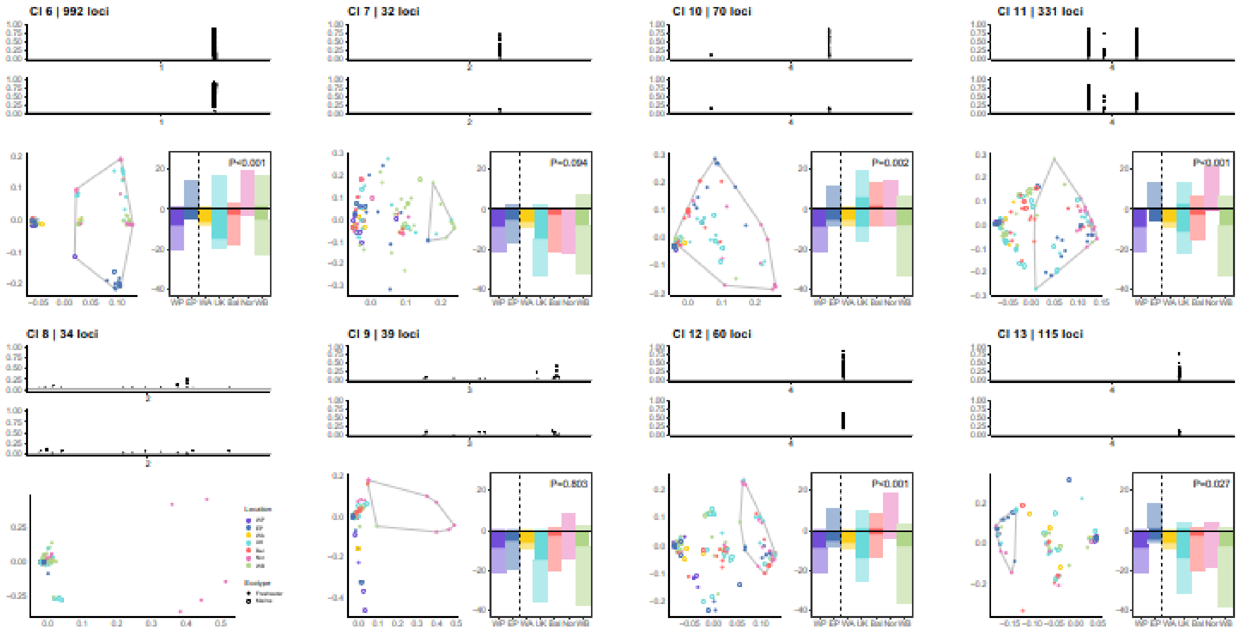
CI 6 | 992 loci
CI 7 | 32 loci
CI 10 | 70 loci
CI 11 | 331 loci
CI 8 | 34 loci
CI 9 | 39 loci
CI 12 | 60 loci
CI 13 | 115 loci

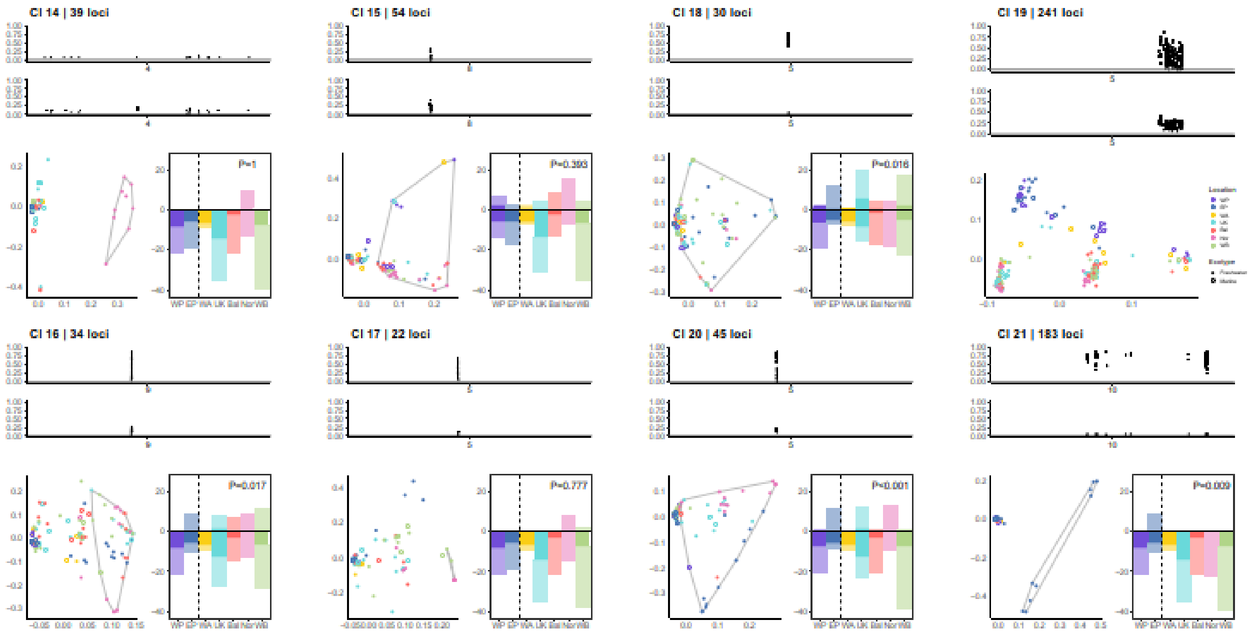

CI 14 | 39 loci
P=1
CI 15 | 54 loci
P=0.393
CI 18 | 30 loci
P=0.016
CI 19 | 241 loci
CI 16 | 34 loci
P=0.017
CI 17 | 22 loci
P=0.777
CI 20 | 45 loci
P=0.001
CI 21 | 183 loci
P=0.009


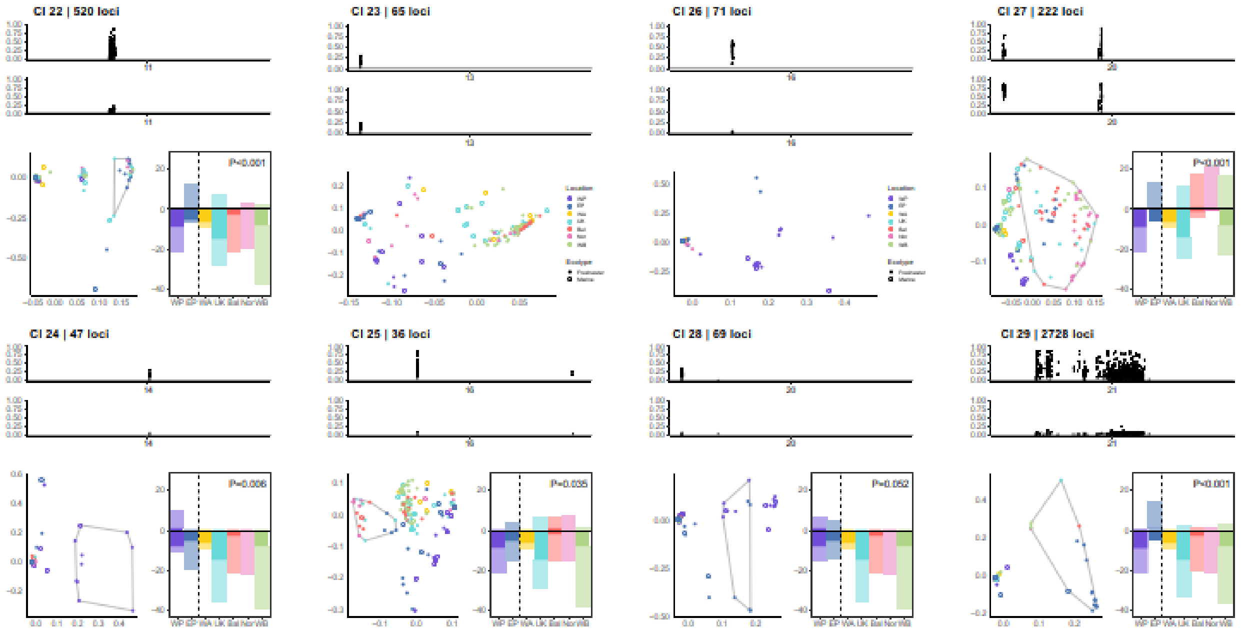

CI 22 | 520 loci
P=0.001
CI 23 | 65 loci
CI 26 | 71 loci
CI 27 | 222 loci
P=0.001
CI 24 | 47 loci
P=0.006
CI 25 | 36 loci
P=0.035
CI 28 | 69 loci
P=0.052
CI 29 | 2728 loci
P=0.001

**Supplementary Figure 3 | Ability of LDna to recover marine-freshwater differentiated regions from Jones et al. (2012).** Jones et al. (2012) identified 812 regions showing parallel marine-freshwater differentiation in the Eastern Pacific ("*i*-regions") and 81 regions showing global parallelism ("*m-f* regions"). (a) The proportions of *m-f* and *i*-regions that were correctly recovered by LDna (red; at least one SNP from 29 LD-clusters mapped to these regions), the proportion or regions for which we had data but LDna analyses failed to recover (cyan), and regions for which we had no genetic data (blue). (b) Number of high LD-SNPs (produced by the first LDna-filtering step) and raw SNPs (bottom row) in regions that were and were not recovered by LDna and (c) size of the regions that were and were not recovered by LDna (on $log_{10}$ scale). (d) $F_{ST}$ from raw SNPs located within regions that were and were not recovered by LDna. Overall, *m-f* regions and *i*- regions that were not recovered by LDna were generally smaller, contained fewer SNPs (i.e. had lower sequencing coverage) and exhibited lower $F_{ST}$ than the regions correctly recovered by LDna.

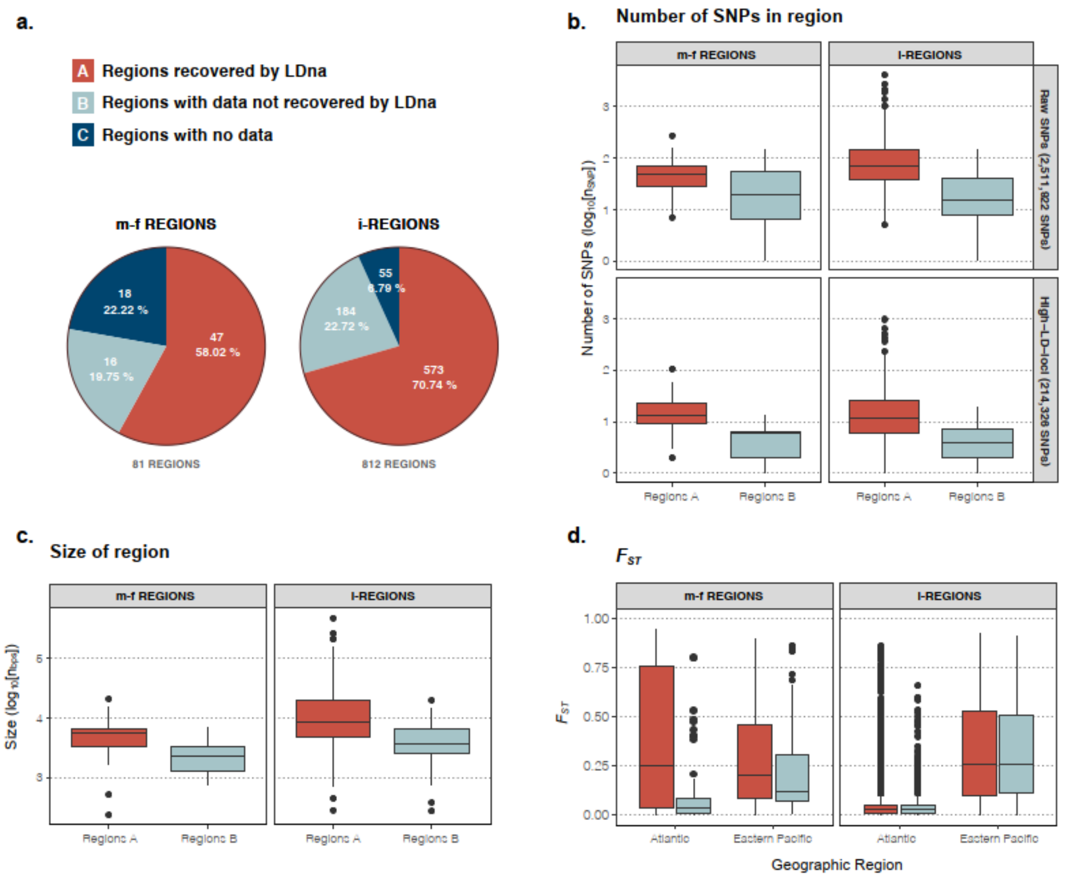

**Supplementary Figure 4 | Genome-wide marine-freshwater differentiation (*$F_{ST}$*) in the Atlantic, Eastern Pacific and Western Pacific Oceans.** (a-c) SNP-based $F_{ST}$ of the individuals in the Atlantic (ATL), Eastern Pacific (EP) and Western Pacific (WP), respectively. Ecotype pairs follow the main analyses (Supplementary Table 2). (d) Window-based $F_{ST}$ (win-size=100 kb) between EP freshwater samples (n=13) and EP marine samples (n=4). (e) Window-based $F_{ST}$ between EP freshwater samples (n=13) and all Pacific marine samples (n=13). (d) and (e) are significantly correlated (*r*=0.904, p<0.0001). (f) and (g) The EP genetic parallelism (LD-clusters 2, 21, 29) shown based on SNP-based $F_{ST}$ under ecotype comparison of (d) and (e) respectively. Loci from LD-clusters involved in genetic parallelism are colour-coded for all panels (refer to main Fig. 2).

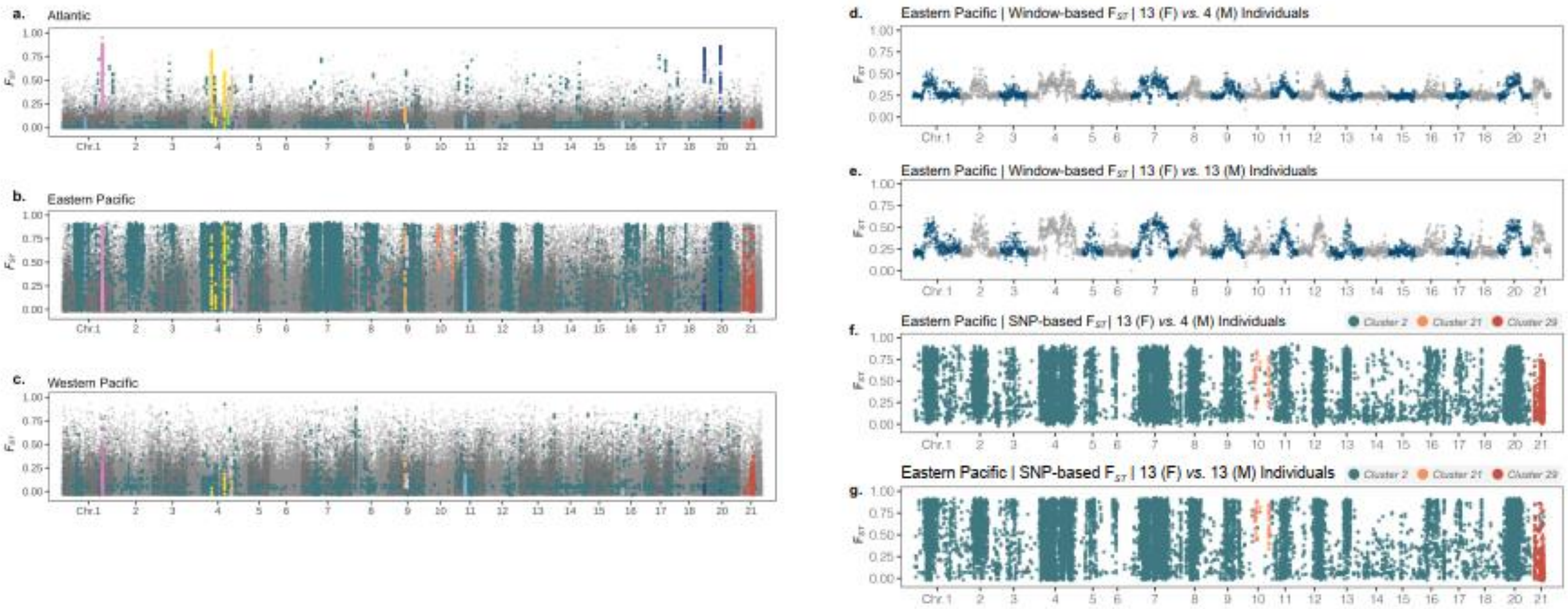

**Supplementary Figure 5 | PCA plot of LDna clusters with population identification.**
See Supplementary Table 1 for population identifications.

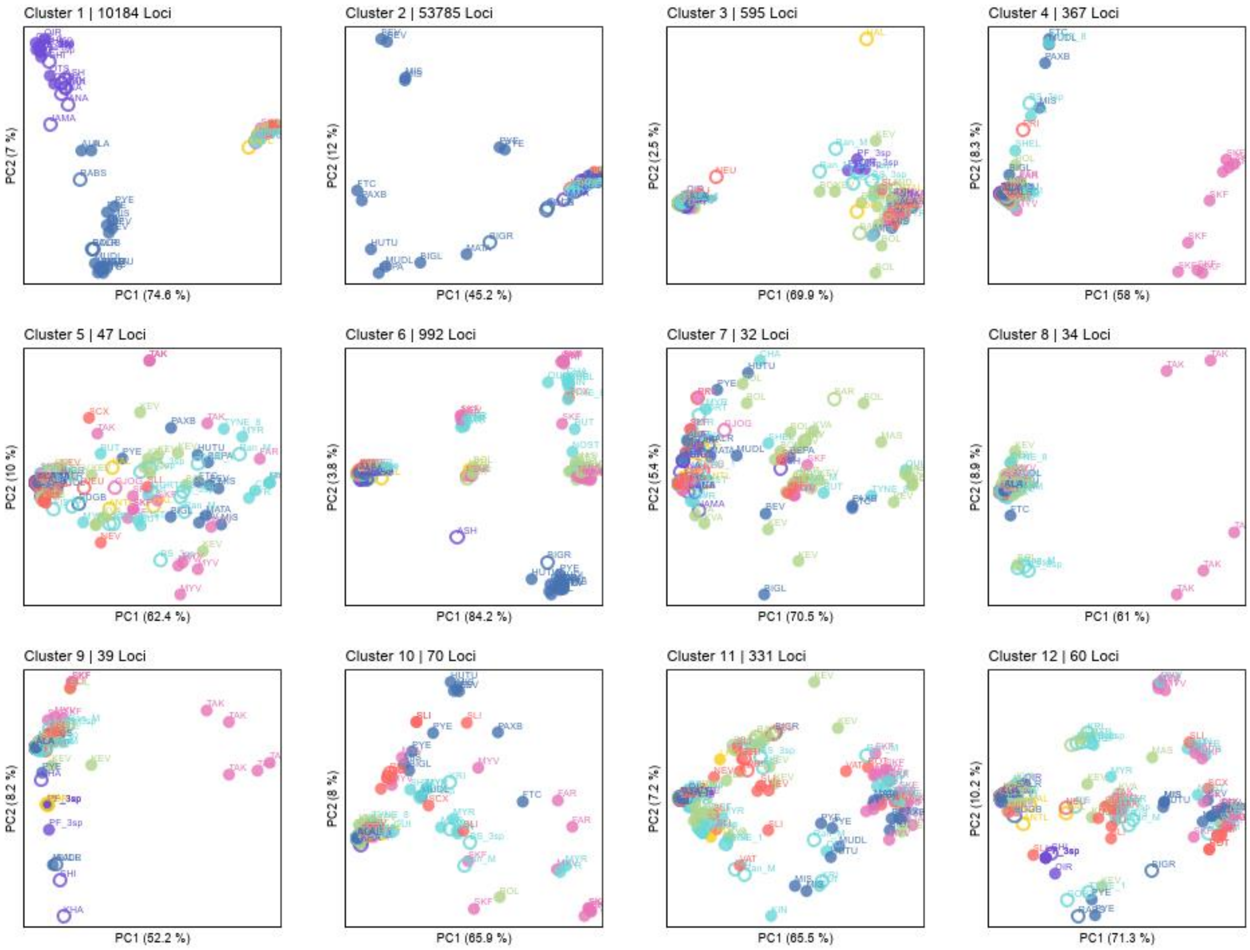

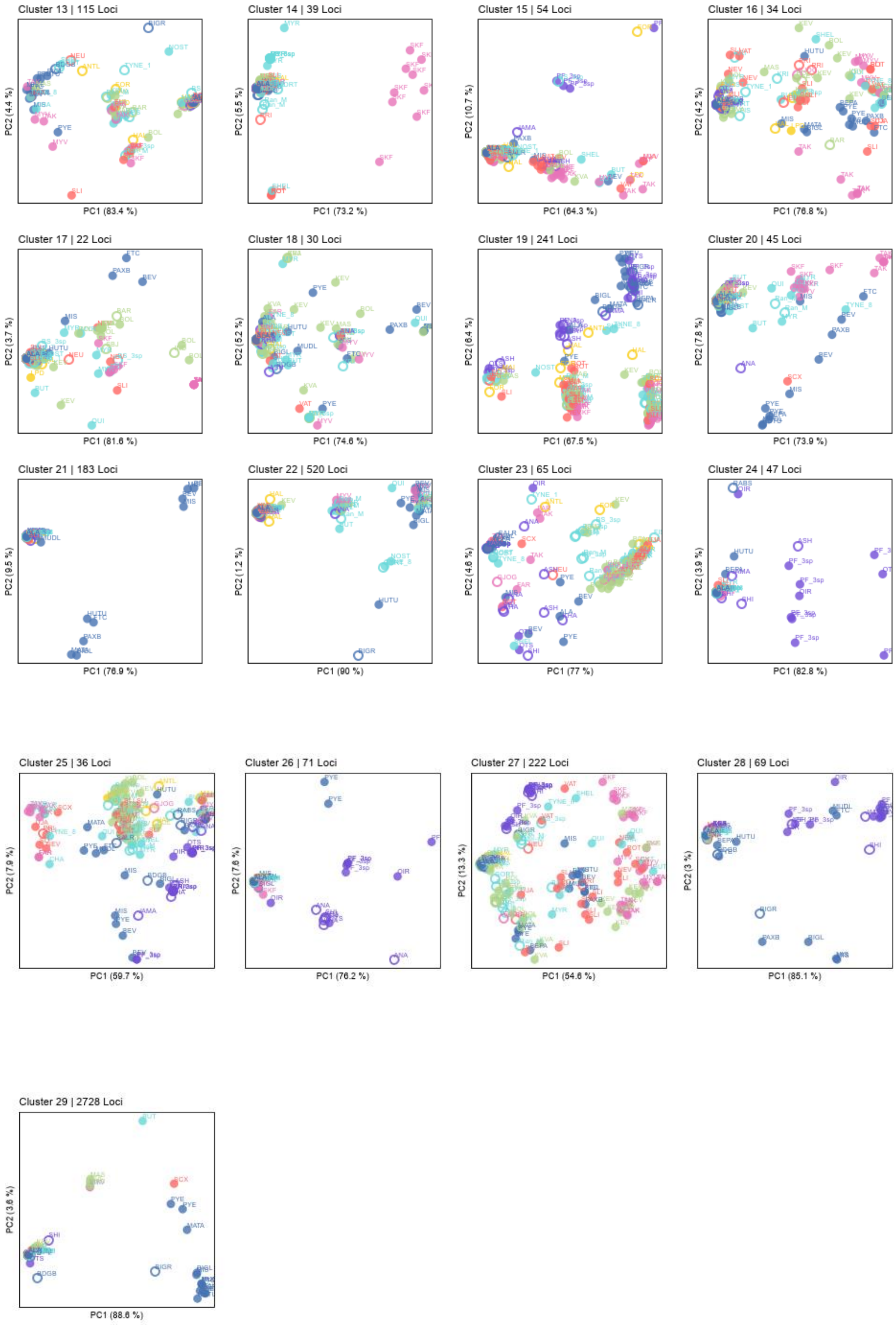
Cluster 13 | 115 Loci
PC2 (4.4 %)
PC1 (83.4 %)
Cluster 14 | 39 Loci
PC2 (5.5 %)
PC1 (73.2 %)
Cluster 15 | 54 Loci
PC2 (10.7 %)
PC1 (64.3 %)
Cluster 16 | 34 Loci
PC2 (4.2 %)
PC1 (76.8 %)
Cluster 17 | 22 Loci
PC2 (3.7 %)
PC1 (81.6 %)
Cluster 18 | 30 Loci
PC2 (5.2 %)
PC1 (74.6 %)
Cluster 19 | 241 Loci
PC2 (6.4 %)
PC1 (67.5 %)
Cluster 20 | 45 Loci
PC2 (7.8 %)
PC1 (73.9 %)
Cluster 21 | 183 Loci
PC2 (9.5 %)
PC1 (76.9 %)
Cluster 22 | 520 Loci
PC2 (1.2 %)
PC1 (90 %)
Cluster 23 | 65 Loci
PC2 (4.6 %)
PC1 (77 %)
Cluster 24 | 47 Loci
PC2 (3.9 %)
PC1 (82.8 %)
Cluster 25 | 36 Loci
PC2 (7.9 %)
PC1 (59.7 %)
Cluster 26 | 71 Loci
PC2 (7.6 %)
PC1 (76.2 %)
Cluster 27 | 222 Loci
PC2 (13.3 %)
PC1 (54.6 %)
Cluster 28 | 69 Loci
PC2 (3 %)
PC1 (85.1 %)
Cluster 29 | 2728 Loci
PC2 (3.6 %)
PC1 (88.6 %)

**Supplementary Figure 6 | Allele frequency changes in simulated data.** (a) Frequency of selected alleles in the marine populations through generations at high and low levels of trans-oceanic gene flow and three different QTL-densities. (b) Frequency of freshwater-adapted alleles in the freshwater populations under six demographic scenarios through generations. (c) Frequency of freshwater-adapted alleles in the marine populations under six demographic scenarios through generations. (d) Effects of recombination rate on marine-freshwater differentiation ($F_{ST}$) of present-day Eastern Pacific and Atlantic populations. The boxplot shows the $F_{ST}$ of loci in high and low recombination genomic regions. (e) Cluster separation score (CSS) of the simulated and empirical data in the Atlantic ocean. For (b) and (c), allele frequency is shown separately for three categories of genomic regions: whole genome (all), high and low recombination genomic regions.

(a)

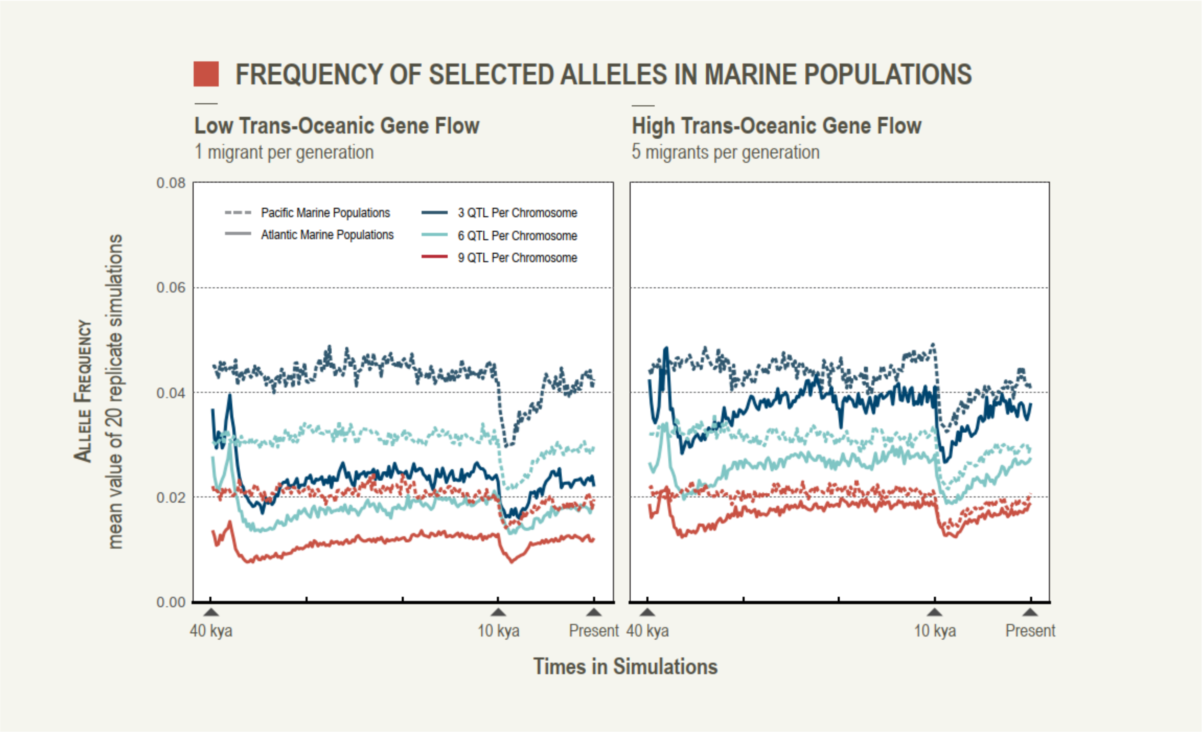

(b)

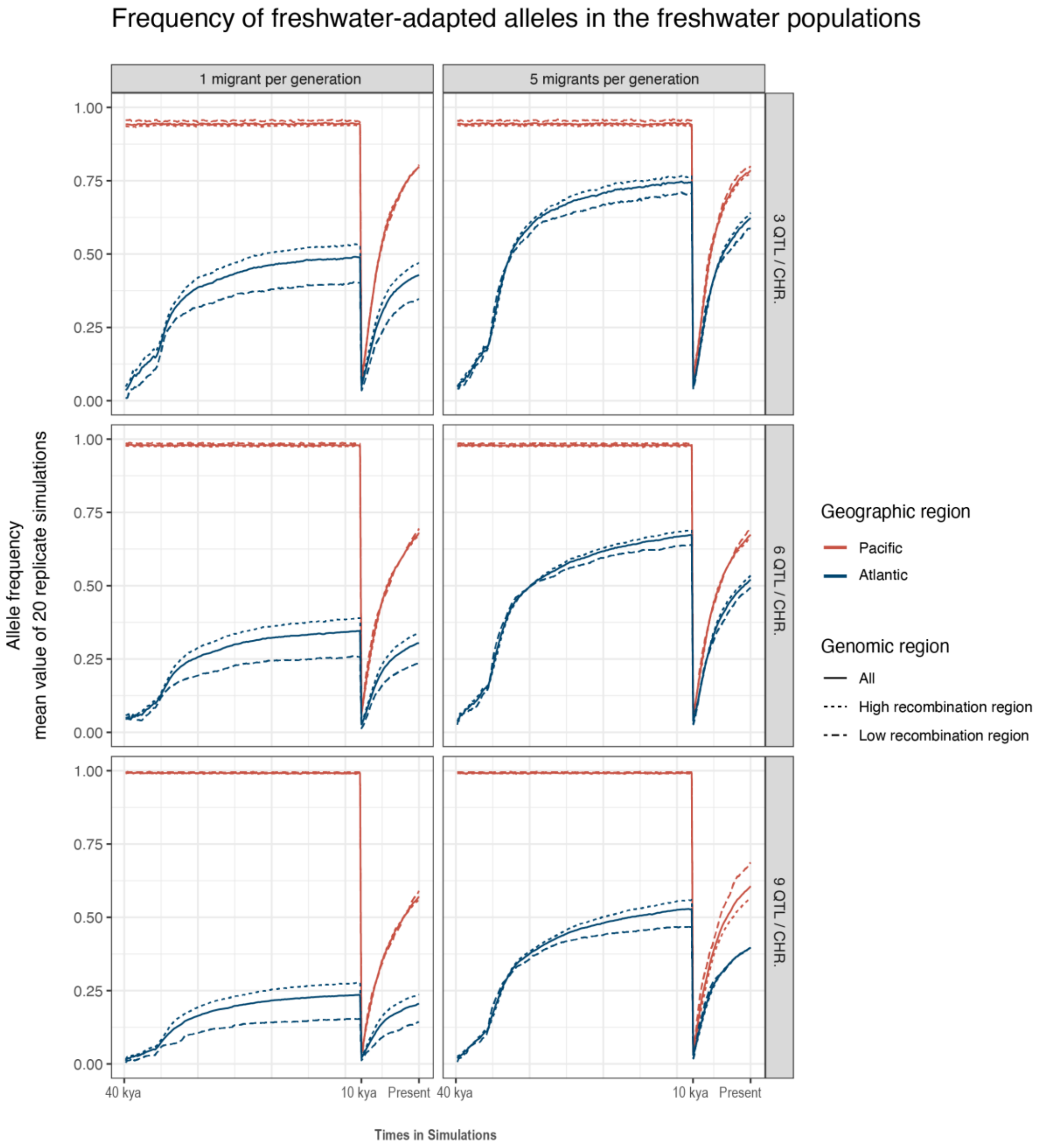

Frequency of freshwater-adapted alleles in the freshwater populations
1 migrant per generation
5 migrants per generation
3 QTL / CHR.
6 QTL / CHR.
9 QTL / CHR.
Allele frequency
mean value of 20 replicate simulations
1.00
0.75
0.50
0.25
0.00
40 kya
10 kya
Present
Times in Simulations
Geographic region
Pacific
Atlantic
Genomic region
All
High recombination region
Low recombination region

(c)

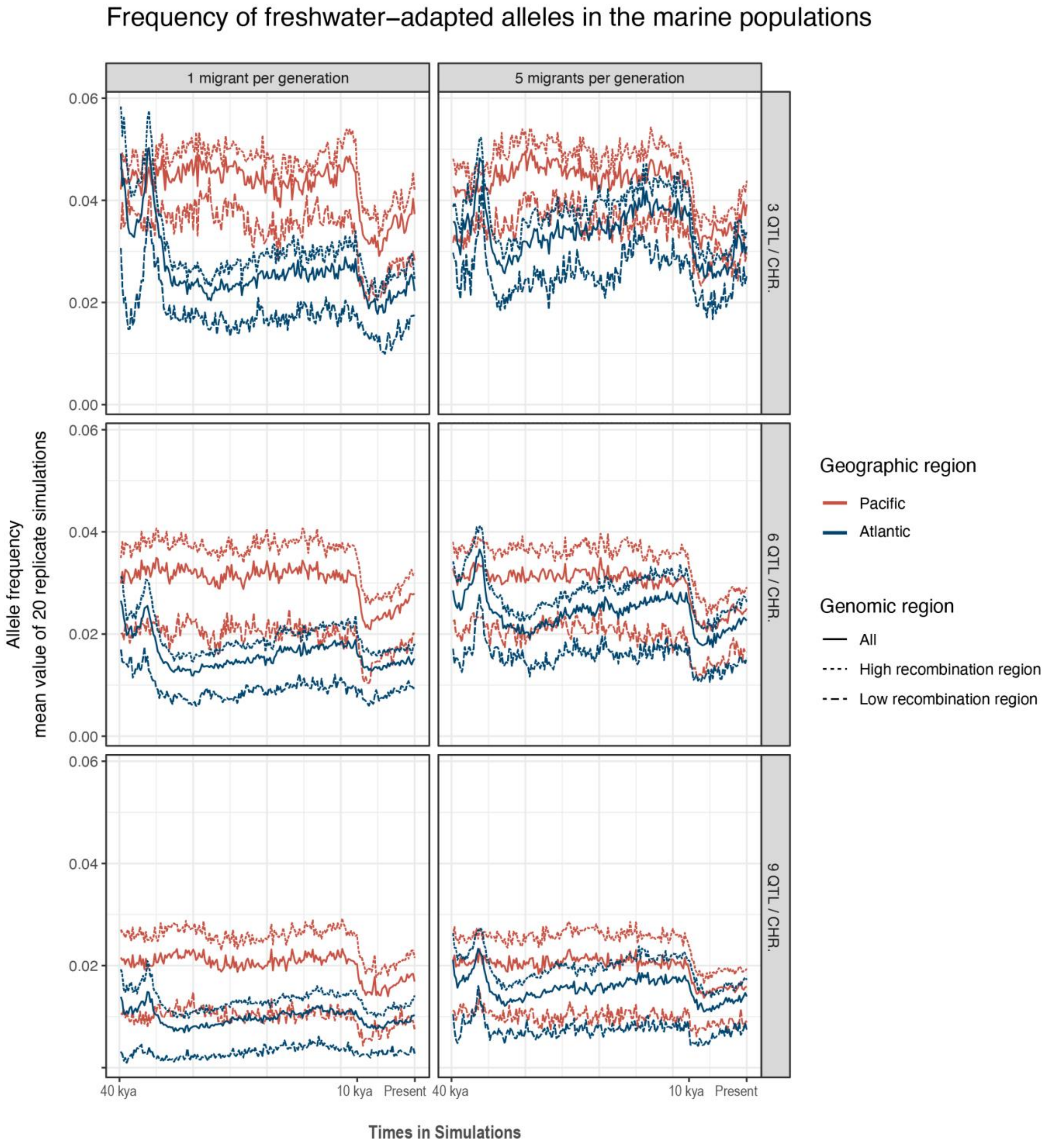

Frequency of freshwater−adapted alleles in the marine populations
1 migrant per generation
5 migrants per generation
3 QTL / CHR.
6 QTL / CHR.
9 QTL / CHR.
Allele frequency
mean value of 20 replicate simulations
0.06
0.04
0.02
0.00
40 kya
10 kya
Present
Times in Simulations
Geographic region
Pacific
Atlantic
Genomic region
All
High recombination region
Low recombination region

(d)

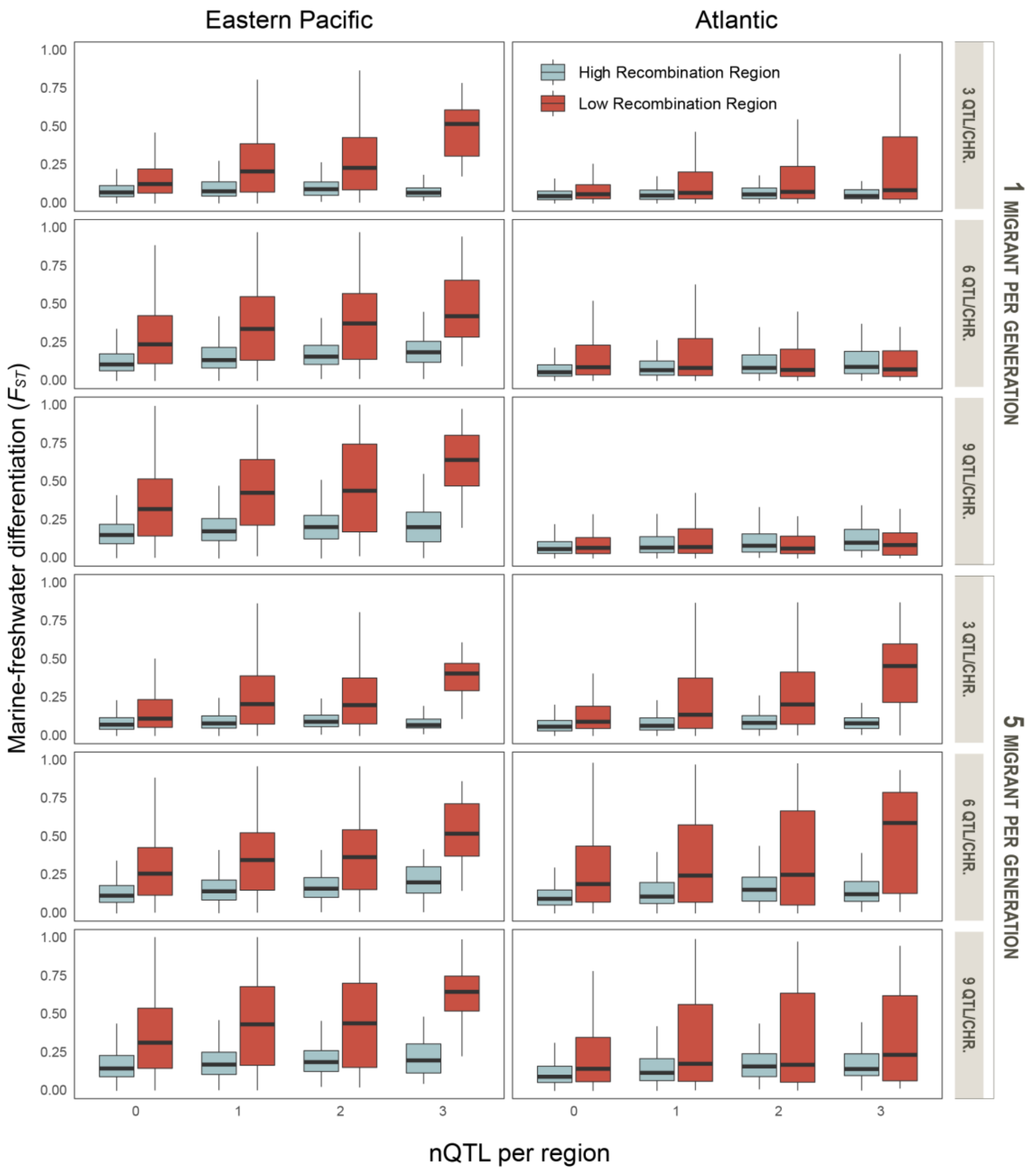

Eastern Pacific
Atlantic
High Recombination Region
Low Recombination Region
Marine-freshwater differentiation ($F_{ST}$)
1.00
0.75
0.50
0.25
0.00
0
1
2
3
nQTL per region
3 QTL/CHR.
6 QTL/CHR.
9 QTL/CHR.
1 MIGRANT PER GENERATION
5 MIGRANT PER GENERATION

**(e)**

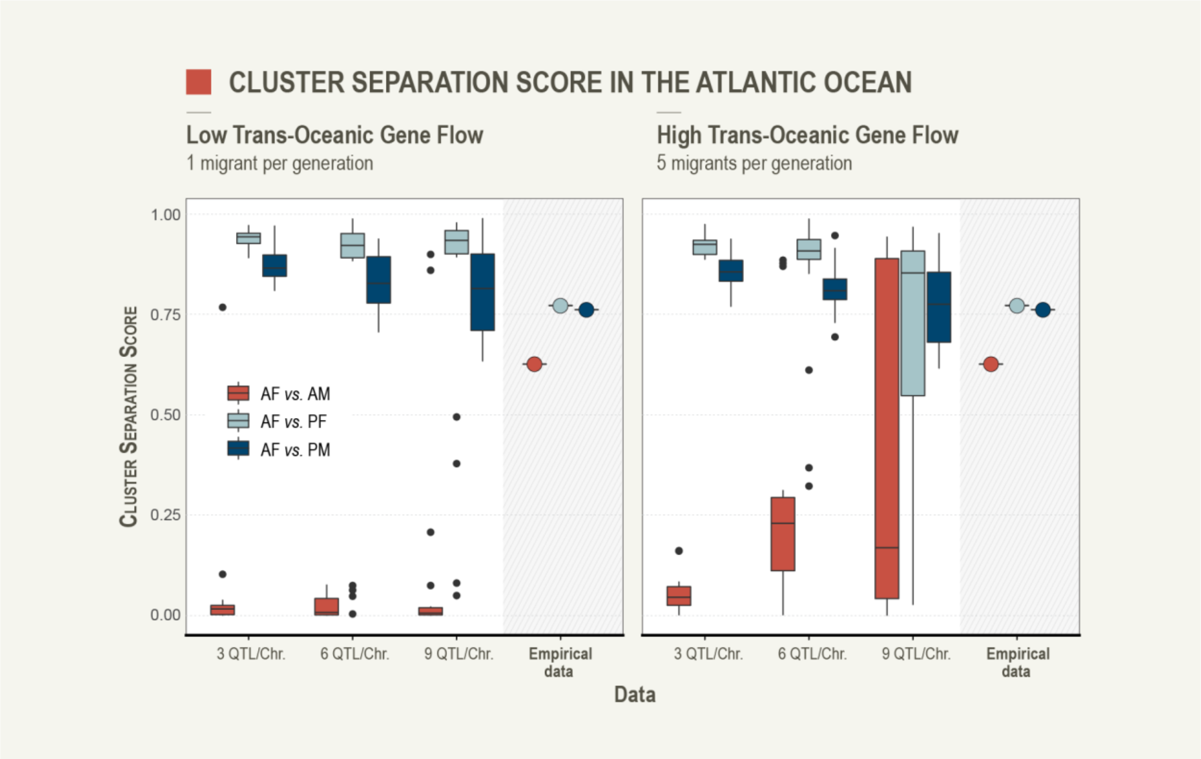

CLUSTER SEPARATION SCORE IN THE ATLANTIC OCEAN
Low Trans-Oceanic Gene Flow
1 migrant per generation
High Trans-Oceanic Gene Flow
5 migrants per generation
Cluster Separation Score
1.00
0.75
0.50
0.25
0.00
AF vs. AM
AF vs. PF
AF vs. PM
3 QTL/Chr.
6 QTL/Chr.
9 QTL/Chr.
Empirical data
Data

**Supplementary Figure 7 | Population variation and Isolation-By-Distance (IBD) in marine three-spined stickleback populations.** (a) Individual heterozygosity (proportion heterozygous loci per individual) of marine individuals in different geographical regions (EP = Eastern Pacific, WP = Western Pacific and ATL = Atlantic). (a) Individual heterozygosity of LD-cluster 2 in different regions. (c) Isolation-by-distance (IBD) between marine populations.

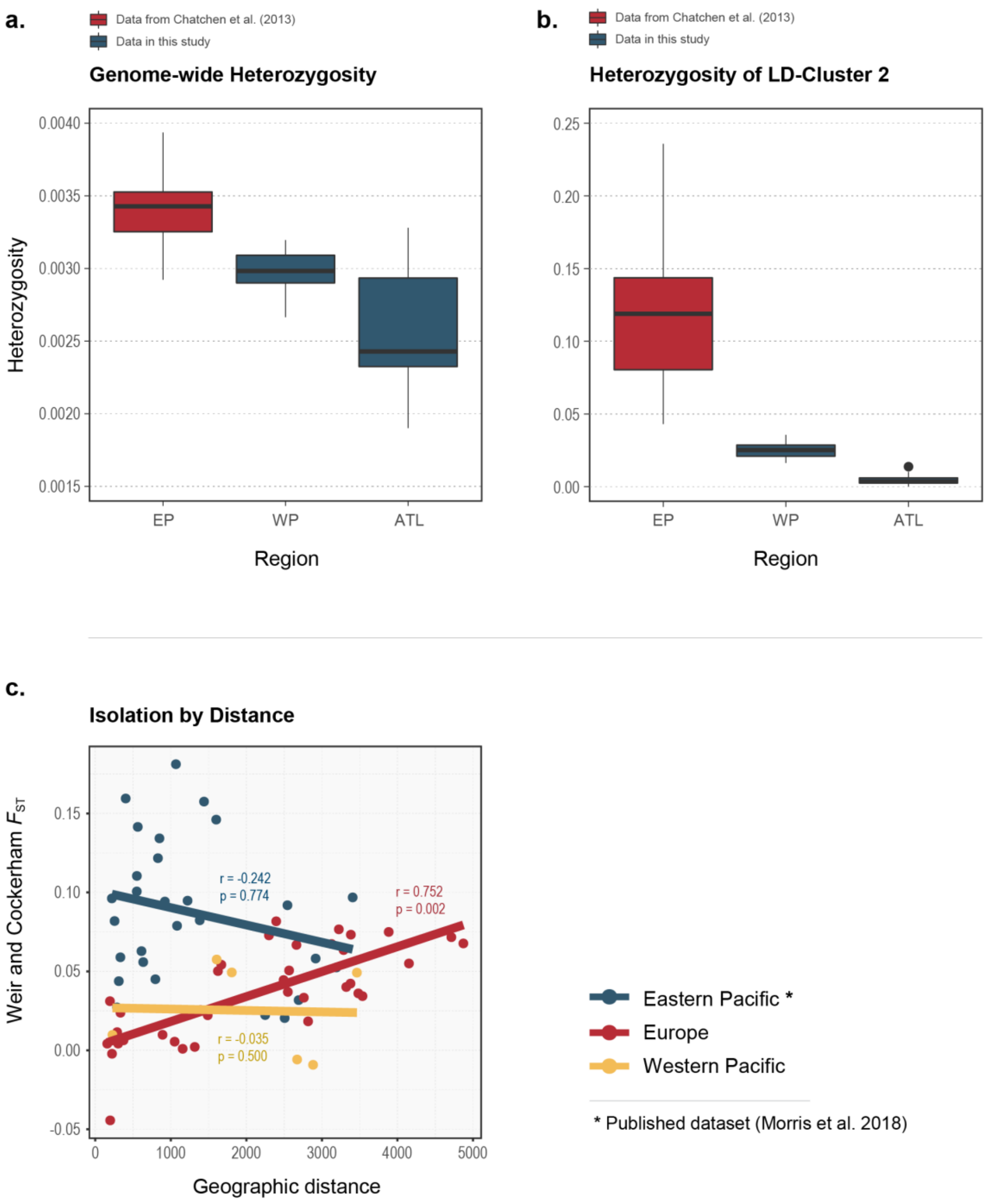

**Supplementary Figure 8 | The 29 clusters identified by LDna in a network tree.** The tree is produced in the final step of LDna (Supplementary Information 1). The 81 tips of the tree indicate the Maximally Connected Loci from each of the clusters identified in the previous steps. The final LDna analyses use these loci to group outlier clusters from the different chromosomes (i.e. the clusters from the second step) into the 29 final LD-clusters used in our analyses.

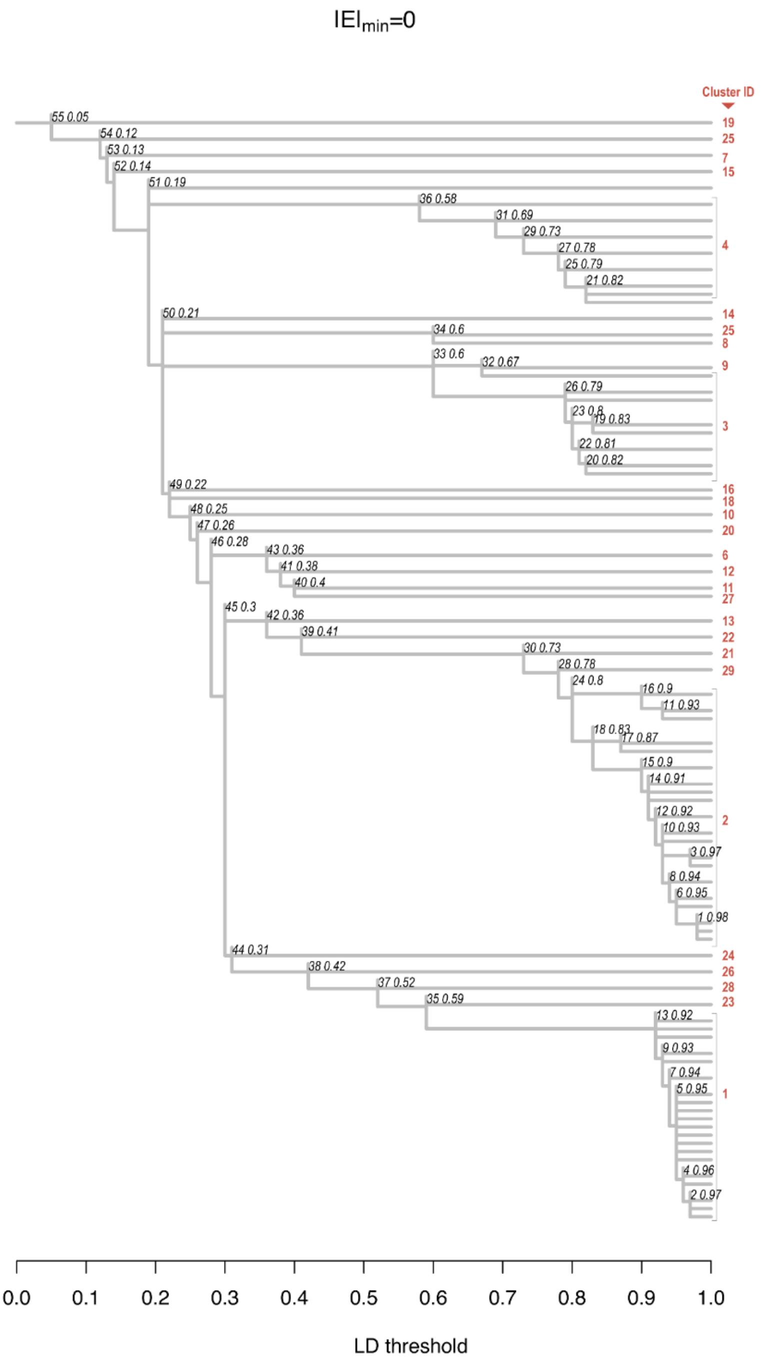

**Supplementary Figure 9 | A chromosome in the simulation.** The plot illustrates the recombination distance (y-axis; centiMorgan, cM) versus physical distance between 100 loci. The recombination distance was shorter at the chromosome centres to simulate the presence of centromeres. The grey shaded area depicts the low recombination region in the chromosome, according to loci position (x-axis).

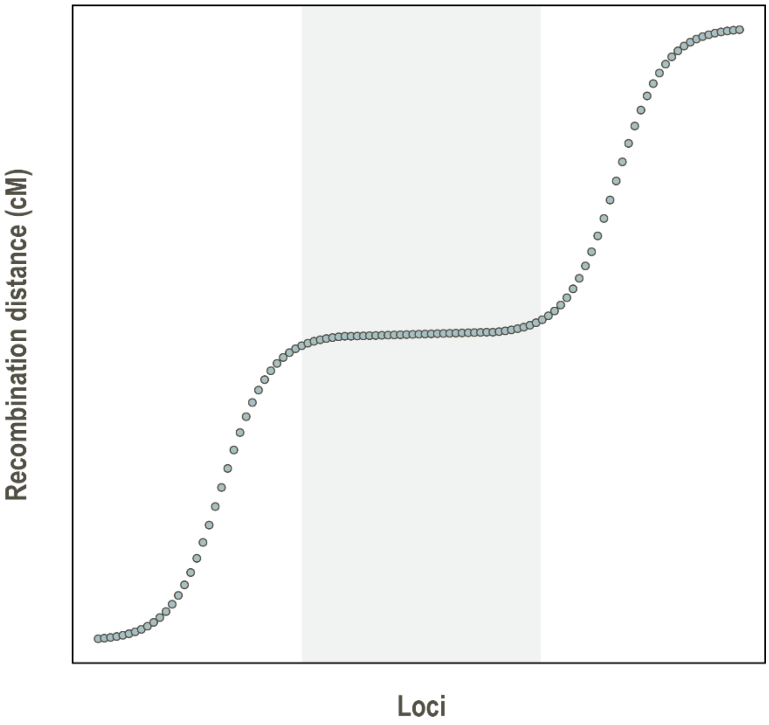

#### Supplementary Table 1 | Details of the samples used in the study.

| Sample ID | Population ID | Ecotype | Region | GPS_N | GPS_E | Coverage | Source of sequences |
|---|---|---|---|---|---|---|---|
| RUS-BOL-GA-1 | BOL | Freshwater | White and Barents Sea | 66.295278 | 33.366111 | 5.79 | This study |
| RUS-BOL-GA-15 | BOL | Freshwater | White and Barents Sea | 66.295278 | 33.366111 | 5.77 | This study |
| RUS-BOL-GA-16 | BOL | Freshwater | White and Barents Sea | 66.295278 | 33.366111 | 5.28 | This study |
| RUS-BOL-GA-17 | BOL | Freshwater | White and Barents Sea | 66.295278 | 33.366111 | 7.40 | Fang et al. [1] |
| RUS-BOL-GA-18 | BOL | Freshwater | White and Barents Sea | 66.295278 | 33.366111 | 4.38 | This study |
| RUS-BOL-GA-20 | BOL | Freshwater | White and Barents Sea | 66.295278 | 33.366111 | 3.07 | This study |
| RUS-BOL-GA-21 | BOL | Freshwater | White and Barents Sea | 66.295278 | 33.366111 | 4.30 | This study |
| RUS-BOL-GA-2 | BOL | Freshwater | White and Barents Sea | 66.295278 | 33.366111 | 5.28 | Fang et al. [1] |
| RUS-BOL-GA-22 | BOL | Freshwater | White and Barents Sea | 66.295278 | 33.366111 | 5.15 | This study |
| RUS-BOL-GA-23 | BOL | Freshwater | White and Barents Sea | 66.295278 | 33.366111 | 5.64 | This study |
| RUS-MAS-GA-10 | MAS | Freshwater | White and Barents Sea | 66.291944 | 33.381667 | 17.91 | Fang et al. [1] |
| RUS-MAS-GA-3 | MAS | Freshwater | White and Barents Sea | 66.291944 | 33.381667 | 3.48 | This study |
| NOR-KVA-GA-14 | KVA | Freshwater | White and Barents Sea | 70.094444 | 28.978611 | 6.08 | This study |
| NOR-KVA-GA-15 | KVA | Freshwater | White and Barents Sea | 70.094444 | 28.978611 | 5.70 | This study |
| NOR-KVA-GA-16 | KVA | Freshwater | White and Barents Sea | 70.094444 | 28.978611 | 10.75 | This study |
| NOR-KVA-GA-19 | KVA | Freshwater | White and Barents Sea | 70.094444 | 28.978611 | 6.27 | This study |
| NOR-KVA-GA-23 | KVA | Freshwater | White and Barents Sea | 70.094444 | 28.978611 | 6.49 | This study |
| NOR-KVA-GA-39 | KVA | Freshwater | White and Barents Sea | 70.094444 | 28.978611 | 10.33 | This study |
| NOR-KVA-GA-42 | KVA | Freshwater | White and Barents Sea | 70.094444 | 28.978611 | 10.64 | Fang et al. [1] |
| NOR-KVA-GA-44 | KVA | Freshwater | White and Barents Sea | 70.094444 | 28.978611 | 7.29 | Fang et al. [1] |
| NOR-KVA-GA-49 | KVA | Freshwater | White and Barents Sea | 70.094444 | 28.978611 | 6.02 | This study |
| NOR-KVA-GA-50 | KVA | Freshwater | White and Barents Sea | 70.094444 | 28.978611 | 4.24 | This study |
| FIN-KEV-GA-48 | KEV | Freshwater | White and Barents Sea | 69.750278 | 27.015278 | 9.85 | Fang et al. [1] |
| FIN-KEV-GA-37 | KEV | Freshwater | White and Barents Sea | 69.750278 | 27.015278 | 8.32 | Fang et al. [1] |
| FIN-KEV-GA-38 | KEV | Freshwater | White and Barents Sea | 69.750278 | 27.015278 | 2.99 | This study |
| FIN-KEV-GA-39 | KEV | Freshwater | White and Barents Sea | 69.750278 | 27.015278 | 1.88 | This study |
| FIN-KEV-GA-41 | KEV | Freshwater | White and Barents Sea | 69.750278 | 27.015278 | 2.65 | This study |
| FIN-KEV-GA-42 | KEV | Freshwater | White and Barents Sea | 69.750278 | 27.015278 | 3.33 | This study |
| FIN-KEV-GA-44 | KEV | Freshwater | White and Barents Sea | 69.750278 | 27.015278 | 6.42 | This study |
| FIN-KEV-GA-46 | KEV | Freshwater | White and Barents Sea | 69.750278 | 27.015278 | 4.80 | This study |
| FIN-KEV-GA-47 | KEV | Freshwater | White and Barents Sea | 69.750278 | 27.015278 | 2.78 | This study |
| GAC-SWE-R-13 | ROT | Freshwater | Baltic Sea | 64.265000 | 20.318889 | 9.93 | Fang et al. [1] |
| GAC-SWE-R-16 | ROT | Freshwater | Baltic Sea | 64.265000 | 20.318889 | 9.02 | Fang et al. [1] |
| rou-A-40 | VAT | Freshwater | Baltic Sea | 58.646111 | 14.638611 | 11.93 | Fang et al. [1] |
| rou-A-41 | VAT | Freshwater | Baltic Sea | 58.646111 | 14.638611 | 11.93 | Fang et al. [1] |
| Rus-Sli-9 | SLI | Freshwater | Baltic Sea | 59.936944 | 31.022778 | 4.87 | This study |
| Rus-Sli-10 | SLI | Freshwater | Baltic Sea | 59.936944 | 31.022778 | 3.88 | This study |
| RUS-SLI-GA-7 | SLI | Freshwater | Baltic Sea | 59.936944 | 31.022778 | 6.32 | This study |
| RUS-SLI-GA-8 | SLI | Freshwater | Baltic Sea | 59.936944 | 31.022778 | 5.19 | This study |
| RUS-SLI-GA-4 | SLI | Freshwater | Baltic Sea | 59.936944 | 31.022778 | 7.16 | Fang et al. [1] |
| RUS-SLI-GA-2 | SLI | Freshwater | Baltic Sea | 59.936944 | 31.022778 | 5.40 | This study |
| RUS-SLI-GA-3 | SLI | Freshwater | Baltic Sea | 59.936944 | 31.022778 | 5.64 | This study |
| RUS-SLI-GA-5 | SLI | Freshwater | Baltic Sea | 59.936944 | 31.022778 | 11.76 | Fang et al. [1] |
| Rus-Sli-6 | SLI | Freshwater | Baltic Sea | 59.936944 | 31.022778 | 7.91 | Fang et al. [1] |
| RUS-SLI-GA-1 | SLI | Freshwater | Baltic Sea | 59.936944 | 31.022778 | 5.14 | This study |
| RUS-UIA-GA-18 | UIA | Freshwater | Baltic Sea | 62.146667 | 35.281389 | 10.53 | Fang et al. [1] |
| LIT-NEV-GA-39 | NEV | Freshwater | Baltic Sea | 54.638889 | 25.364167 | 7.77 | Fang et al. [1] |
| LIT-NEV-GA-6 | NEV | Freshwater | Baltic Sea | 54.638889 | 25.364168 | 8.88 | Fang et al. [1] |
| SCX | SCX | Freshwater | Baltic Sea | 54.080000 | 10.083000 | 1.83 | Jones et al. [2] |
| NOR-SKF-GA-43 | SKF | Freshwater | Norwegian Sea | 69.956667 | 19.145833 | 8.23 | This study |
| NOR-SKF-GA-41 | SKF | Freshwater | Norwegian Sea | 69.956667 | 19.145833 | 13.39 | This study |
| NOR-SKF-GA-46 | SKF | Freshwater | Norwegian Sea | 69.956667 | 19.145833 | 8.48 | This study |
| NOR-SKF-GA-45 | SKF | Freshwater | Norwegian Sea | 69.956667 | 19.145833 | 9.65 | This study |
| NOR-SKF-GA-6 | SKF | Freshwater | Norwegian Sea | 69.956667 | 19.145833 | 7.99 | This study |
| NOR-SKF-GA-38 | SKF | Freshwater | Norwegian Sea | 69.956667 | 19.145833 | 8.96 | This study |
| NOR-SKF-GA-22 | SKF | Freshwater | Norwegian Sea | 69.956667 | 19.145833 | 3.02 | This study |
| NOR-SKF-GA-33 | SKF | Freshwater | Norwegian Sea | 69.956667 | 19.145833 | 11.34 | Fang et al. [1] |
| NOR-SKF-GA-30 | SKF | Freshwater | Norwegian Sea | 69.956667 | 19.145833 | 10.41 | Fang et al. [1] |
| NOR-TAK-GA-2 | TAK | Freshwater | Norwegian Sea | 69.117222 | 19.067500 | 15.61 | This study |
| NOR-TAK-GA-3 | TAK | Freshwater | Norwegian Sea | 69.117222 | 19.067500 | 17.48 | This study |
| NOR-TAK-GA-5 | TAK | Freshwater | Norwegian Sea | 69.117222 | 19.067500 | 13.06 | Fang et al. [1] |
| NOR-TAK-GA-4 | TAK | Freshwater | Norwegian Sea | 69.117222 | 19.067500 | 6.31 | Fang et al. [1] |
| NOR-TAK-GA-1 | TAK | Freshwater | Norwegian Sea | 69.117222 | 19.067501 | 7.44 | This study |
| F-zhi-KLU-GA-1 | FAR | Freshwater | Norwegian Sea | 62.150278 | -6.633889 | 6.37 | Fang et al. [1] |
| F-zhi-KLU-GA-2 | FAR | Freshwater | Norwegian Sea | 62.150278 | -6.633889 | 29.21 | Fang et al. [1] |
| MYV-M5 | MYV | Freshwater | Norwegian Sea | 65.642222 | -16.927222 | 11.24 | Fang et al. [1] |
| MYV-M2 | MYV | Freshwater | Norwegian Sea | 65.642222 | -16.927222 | 5.09 | This study |
| MYV-M3 | MYV | Freshwater | Norwegian Sea | 65.642222 | -16.927222 | 19.43 | This study |
| MYV-M4 | MYV | Freshwater | Norwegian Sea | 65.642222 | -16.927222 | 11.11 | This study |
| MYV-M1 | MYV | Freshwater | Norwegian Sea | 65.642222 | -16.927222 | 3.15 | This study |
| NOR-MYR-GA-44 | MYR | Freshwater | North Sea and British Isles | 60.313333 | 5.399167 | 17.71 | This study |
| NOR-MYR-GA-42 | MYR | Freshwater | North Sea and British Isles | 60.313333 | 5.399167 | 9.55 | Fang et al. [1] |
| NOR-MYR-GA-3 | MYR | Freshwater | North Sea and British Isles | 60.313333 | 5.399167 | 4.95 | This study |
| NOR-MYR-GA-13 | MYR | Freshwater | North Sea and British Isles | 60.313333 | 5.399167 | 3.60 | This study |
| Nor-Myr-4 | MYR | Freshwater | North Sea and British Isles | 60.313333 | 5.399167 | 10.58 | This study |
| NOR-MYR-GA-47 | MYR | Freshwater | North Sea and British Isles | 60.313333 | 5.399167 | 4.48 | This study |
| NOR-MYR-GA-11 | MYR | Freshwater | North Sea and British Isles | 60.313333 | 5.399167 | 3.71 | This study |
| NOR-MYR-GA-14 | MYR | Freshwater | North Sea and British Isles | 60.313333 | 5.399167 | 9.94 | This study |
| NOR-MYR-GA-20 | MYR | Freshwater | North Sea and British Isles | 60.313333 | 5.399167 | 14.74 | Fang et al. [1] |

| | | | | | | | |
|---|---|---|---|---|---|---|---|
| NOR-MYR-GA-36 | MYR | Freshwater | North Sea and British Isles | 60.313333 | 5.399167 | 11.78 | This study |
| GA-UK-KIN-4 | KIN | Freshwater | North Sea and British Isles | 56.335000 | -2.789722 | 17.44 | Fang et al. [1] |
| GA-UK-KIN-3 | KIN | Freshwater | North Sea and British Isles | 56.335000 | -2.789722 | 16.95 | Fang et al. [1] |
| BUTE-QUIEN-27 | QUI | Freshwater | North Sea and British Isles | 55.789167 | -5.088611 | 4.83 | Fang et al. [1] |
| BUTE-QUIEN-26 | QUI | Freshwater | North Sea and British Isles | 55.789167 | -5.088611 | 13.55 | Fang et al. [1] |
| ENG-BUT-GA-1 | BUT | Freshwater | North Sea and British Isles | 51.728611 | 0.160556 | 3.75 | This study |
| ENG-BUT-GA-33 | BUT | Freshwater | North Sea and British Isles | 51.728611 | 0.160556 | 4.51 | Fang et al. [1] |
| RAN-CHA-GA-1 | CHA | Freshwater | North Sea and British Isles | 47.445556 | 3.650000 | 16.21 | Fang et al. [1] |
| NOST | NOST | Freshwater | North Sea and British Isles | 59.765000 | 5.712000 | 1.37 | Jones et al. [2] |
| TYNE_8 | TYNE_8 | Freshwater | North Sea and British Isles | 55.943000 | -2.785000 | 2.66 | Jones et al. [2] |
| SHEL | SHEL | Freshwater | North Sea and British Isles | 56.748000 | -5.698000 | 2.20 | Jones et al. [2] |
| GA-CA-LPD-2 | LPD | Freshwater | Western Atlantic | 44.577222 | -63.577500 | 10.38 | Fang et al. [1] |
| GA-CA-LPD-3 | LPD | Freshwater | Western Atlantic | 44.577222 | -63.577500 | 10.11 | Fang et al. [1] |
| GA-JP-PO-OIR-4 | OIR | Freshwater | Western Pacific | 40.592778 | 141.222222 | 2.57 | This study |
| GA-JP-PO-OIR-3 | OIR | Freshwater | Western Pacific | 40.592778 | 141.222222 | 5.91 | Fang et al. [1] |
| JP-PO-OTS-GA-10 | OTS | Freshwater | Western Pacific | 39.366667 | 141.883333 | 14.68 | Fang et al. [1] |
| JP-PO-OTS-GA-16 | OTS | Freshwater | Western Pacific | 39.366667 | 141.883333 | 12.64 | Fang et al. [1] |
| PF_JR42 | PF_3sp | Freshwater | Western Pacific | 43.218583 | 144.779694 | 40.81 | Yoshida et al. [3] |
| PF_JR58 | PF_3sp | Freshwater | Western Pacific | 43.218583 | 144.779694 | 18.40 | Yoshida et al. [3] |
| PF_JR65 | PF_3sp | Freshwater | Western Pacific | 43.218583 | 144.779694 | 50.81 | Yoshida et al. [3] |
| PF_JR66 | PF_3sp | Freshwater | Western Pacific | 43.218583 | 144.779694 | 77.19 | Yoshida et al. [3] |
| PF_JR27 | PF_3sp | Freshwater | Western Pacific | 43.218583 | 144.779694 | 14.40 | Yoshida et al. [3] |
| PM_JR39 | PF_3sp | Freshwater | Western Pacific | 43.218583 | 144.779694 | 28.75 | Yoshida et al. [3] |
| PM_JR49 | PF_3sp | Freshwater | Western Pacific | 43.218583 | 144.779694 | 33.09 | Yoshida et al. [3] |
| PM_JR55 | PF_3sp | Freshwater | Western Pacific | 43.218583 | 144.779694 | 40.18 | Yoshida et al. [3] |
| CAN-PYE-GA-6 | PYE | Freshwater | Eastern Pacific | 50.296389 | -125.582222 | 6.58 | Fang et al. [1] |
| CAN-PYE-GA-5 | PYE | Freshwater | Eastern Pacific | 50.296389 | -125.582222 | 3.28 | This study |
| CAN-BEV-GA-3 | BEV | Freshwater | Eastern Pacific | 48.542778 | -123.542222 | 11.84 | Fang et al. [1] |
| CAN-BEV-GA-5 | BEV | Freshwater | Eastern Pacific | 48.542778 | -123.542222 | 7.05 | Fang et al. [1] |
| CAN-MIS-GA-9 | MIS | Freshwater | Eastern Pacific | 53.269444 | -132.067778 | 7.43 | Fang et al. [1] |
| CAN-MIS-GA-4 | MIS | Freshwater | Eastern Pacific | 53.269444 | -132.067778 | 6.04 | Fang et al. [1] |
| USA-ala-GA-23 | ALA | Freshwater | Eastern Pacific | 57.783333 | -152.383333 | 3.67 | This study |
| USA-ala-GA-26 | ALA | Freshwater | Eastern Pacific | 57.783333 | -152.383333 | 20.06 | Fang et al. [1] |
| BEPA | BEPA | Freshwater | Eastern Pacific | 61.615000 | -149.757000 | 1.82 | Jones et al. [2] |
| BIGL | BIGL | Freshwater | Eastern Pacific | 39.3 | -123.728 | 2.84 | Jones et al. [2] |
| FTC | FTC | Freshwater | Eastern Pacific | 48.931 | -122.487 | 2.95 | Jones et al. [2] |
| HUTU | HUTU | Freshwater | Eastern Pacific | 47.231 | -123.957 | 3.41 | Jones et al. [2] |
| MATA | MATA | Freshwater | Eastern Pacific | 37.393 | -122.162 | 2.76 | Jones et al. [2] |
| MUDL | MUDL | Freshwater | Eastern Pacific | 61.593 | -149.34 | 2.03 | Jones et al. [2] |
| PAXB | PAXB | Freshwater | Eastern Pacific | 49.703 | -124.522 | 2.37 | Jones et al. [2] |
| NEU | NEU | Marine | Baltic Sea | 54.058 | 10.877 | 1.34 | Jones et al. [2] |
| GAC-Rus-PRI-12 | PRI | Marine | Baltic Sea | 60.350000 | 28.616667 | 9.81 | Fang et al. [1] |
| GAC-RUS-PRI-33 | PRI | Marine | Baltic Sea | 60.350000 | 28.616667 | 16.15 | Fang et al. [1] |
| NOR-BAR-GA-42 | BAR | Marine | White and Barents Sea | 74.966944 | 37.134167 | 12.13 | Fang et al. [1] |
| NOR-BAR-GA-43 | BAR | Marine | White and Barents Sea | 74.966944 | 37.134167 | 3.27 | This study |
| NOR-SBJ-GA-1 | SBJ | Marine | White and Barents Sea | 71.823333 | 33.043333 | 16.77 | Fang et al. [1] |
| NOR-SBJ-GA-2 | SBJ | Marine | White and Barents Sea | 71.823333 | 33.043333 | 15.92 | Fang et al. [1] |
| RUS-IND-GA-5 | IND | Marine | White and Barents Sea | 66.240000 | 37.145000 | 23.52 | Fang et al. [1] |
| RUS-IND-GA-4 | IND | Marine | White and Barents Sea | 66.240000 | 37.145000 | 18.19 | Fang et al. [1] |
| RUS-LEV-GA-1 | LEV | Marine | White and Barents Sea | 66.290556 | 33.434167 | 10.40 | Fang et al. [1] |
| RUS-LEV-GA-2 | LEV | Marine | White and Barents Sea | 66.290556 | 33.434167 | 14.44 | Fang et al. [1] |
| GJOG | GJOG | Marine | Norwegian Sea | 65.98 | -21.44 | 2.41 | Jones et al. [2] |
| GAC-NOR-KRI-12 | KRI | Marine | North Sea and British Isles | 58.165556 | 8.031111 | 6.24 | Fang et al. [1] |
| GAC-NOR-KRI-35 | KRI | Marine | North Sea and British Isles | 58.165556 | 8.031111 | 9.59 | Fang et al. [1] |
| GAC-SWE-FIS-26 | FIS | Marine | North Sea and British Isles | 58.234722 | 11.401667 | 13.69 | Fang et al. [1] |
| GAC-SWE-FIS-28 | FIS | Marine | North Sea and British Isles | 58.234722 | 11.401667 | 3.06 | This study |
| bsi25 | BS_3sp | Marine | North Sea and British Isles | 56.602553 | 8.300539 | 24.97 | Feulner et al. [4] |
| bsi26 | BS_3sp | Marine | North Sea and British Isles | 56.602553 | 8.300539 | 29.27 | Feulner et al. [4] |
| bsi27 | BS_3sp | Marine | North Sea and British Isles | 56.602553 | 8.300539 | 27.69 | Feulner et al. [4] |
| bsi28 | BS_3sp | Marine | North Sea and British Isles | 56.602553 | 8.300539 | 20.96 | Feulner et al. [4] |
| bsi29 | BS_3sp | Marine | North Sea and British Isles | 56.602553 | 8.300539 | 21.26 | Feulner et al. [4] |
| bsi30 | BS_3sp | Marine | North Sea and British Isles | 56.602553 | 8.300539 | 31.42 | Feulner et al. [4] |
| GORT | GORT | Marine | North Sea and British Isles | 56.911 | -5.888 | 2.29 | Jones et al. [2] |
| TYNE_1 | TYNE_1 | Marine | North Sea and British Isles | 55.999 | -2.52 | 1.72 | Jones et al. [2] |
| GA-US-MAI-1 | MAI | Marine | Western Atlantic | 44.383056 | -68.942222 | 4.22 | This study |
| GA-US-MAI-3 | MAI | Marine | Western Atlantic | 44.383056 | -68.942222 | 19.53 | Fang et al. [1] |
| GA-FOR-M-45 | FOR | Marine | Western Atlantic | 47.717500 | -69.740000 | 5.42 | Fang et al. [1] |
| GA-FOR-F52 | FOR | Marine | Western Atlantic | 47.717500 | -69.740000 | 5.49 | Fang et al. [1] |
| Can-Hal-1 | HAL | Marine | Western Atlantic | 44.647222 | -63.416667 | 1.82 | This study |
| Can-Hal-6 | HAL | Marine | Western Atlantic | 44.647222 | -63.416667 | 2.06 | This study |
| ANTL | ANTL | Marine | Western Atlantic | 45.698 | -61.878 | 1.51 | Jones et al. [2] |
| Ran_M1 | Ran_M | Marine | North Sea and British Isles | 56.606686 | 10.302019 | 5.80 | Liu et al. [5] |
| Ran_M3 | Ran_M | Marine | North Sea and British Isles | 56.606687 | 10.302020 | 5.89 | Liu et al. [5] |
| Ran_M5 | Ran_M | Marine | North Sea and British Isles | 56.606688 | 10.302021 | 5.92 | Liu et al. [5] |
| RUS-AN-GA-10 | ANA | Marine | Western Pacific | 64.735833 | 177.526944 | 17.95 | Fang et al. [1] |
| RUS-AN-GA-2 | ANA | Marine | Western Pacific | 64.735833 | 177.526944 | 7.85 | Fang et al. [1] |
| RUS-KHA-GA-6 | KHA | Marine | Western Pacific | 57.070556 | 156.601667 | 10.73 | Fang et al. [1] |
| RUS-KHA-GA-4 | KHA | Marine | Western Pacific | 57.070556 | 156.601667 | 6.16 | Fang et al. [1] |
| RUS-ASH-GA-3 | ASH | Marine | Western Pacific | 55.312778 | 155.567222 | 7.85 | Fang et al. [1] |
| RUS-ASH-GA-1 | ASH | Marine | Western Pacific | 55.312778 | 155.567222 | 10.67 | Fang et al. [1] |
| GA-PO-SHI-3 | SHI | Marine | Western Pacific | 43.031111 | 144.845556 | 6.31 | Fang et al. [1] |
| GA-PO-SHI-2 | SHI | Marine | Western Pacific | 43.031111 | 144.845556 | 14.55 | Fang et al. [1] |
| JAMA | JAMA | Marine | Western Pacific | 42.973 | 144.329 | 2.62 | Jones et al. [2] |
| BDGB | BDGB | Marine | Eastern Pacific | 38.325 | -123.041 | 2.06 | Jones et al. [2] |
| BIGR | BIGR | Marine | Eastern Pacific | 39.289 | -123.747 | 2.14 | Jones et al. [2] |
| RABS | RABS | Marine | Eastern Pacific | 61.556 | -149.249 | 2.99 | Jones et al. [2] |
| SALR | SALR | Marine | Eastern Pacific | 49.175 | -122.594 | 1.83 | Jones et al. [2] |


1 Fang, B., Merila, J., Ribeiro, F., Alexandre, C. M. & Momigliano, P. Worldwide phylogeny of three-spined sticklebacks. *Mol. Phylogenet. Evol.* **127**, 613-625, doi:10.1016/j.ympev.2018.06.008 (**2018**).

2 Jones, F. C. *et al.* The genomic basis of adaptive evolution in threespine sticklebacks. *Nature* **484**, 55-61, doi:10.1038/nature10944 (**2012**).

3 Yoshida, K. *et al.* Sex chromosome turnover contributes to genomic divergence between incipient stickleback species. *PLoS Genet.* **10**, e1004223, doi:10.1371/journal.pgen.1004223 (**2014**).

4 Feulner, P. G. *et al.* Genome-wide patterns of standing genetic variation in a marine population of three-spined sticklebacks. *Mol. Ecol.* **22**, 635-649, doi:10.1111/j.1365-294X.2012.05680.x (**2013**).

5 Liu, S., Hansen, M. M. & Jacobsen, M. W. Region-wide and ecotype-specific differences in demographic histories of threespine stickleback populations, estimated from whole genome sequences. *Mol. Ecol.* **25**, 5187-5202, doi:10.1111/mec.13827 (**2016**).

**Supplementary Table 2 | Specification of different sampling schemes for the $F_{ST}$ analyses between marine and freshwater populations.**

| No. | Figrue ID | Region | Freshwater samples | Marine samples |
|---|---|---|---|---|
| 1 | Fig. 2b,e; Suppl. Fig. 2b,e,g | Eastern Pacific (EP) | 13 EP | 4 EP + 9 WP |
| 2 | Fig. 2d; Suppl. Fig. 2a | Atlantic (ATL) | 92 ATL | 34 ATL |
| 3 | Suppl. Fig. 4c | Western Pacific (WP) | 12 WP | 4 EP + 9 WP |

**Supplementary Table 3 | Summary of all LD-clusters.** Shaded rows (LD-clusters) contribute to genetic parallelism of regional or trans-oceanic freshwater populations.

| Cluster ID | $N_{loci}$ | $LD_{mean}$ | $LD_{median}$ | $P_{randomisation\text{-}test}$ | Located Chromosome | $N_{overlap\_ecotype\_divergent\_regions}$ (regions/SNPs) * | $N_{overlap\_tree\_i}$ (regions/SNPs) ** | Inferred cause |
|---|---|---|---|---|---|---|---|---|
| Cluster 1 | 10184 | 0.536 | 0.537 | NA | All chromosomes | 1/2 | 50/153 | Geographic structure |
| Cluster 2 | 53790 | 0.385 | 0.391 | 0.001 | All chromosomes | 12/28 | 539/8482 | Local adaptation (genetic parallelism of **Eastern Pacific** freshwater populations) |
| Cluster 3 | 595 | 0.395 | 0.398 | 0.252 | Chr. I, IV, VI, VIII, IX, XI, XII, XIV, XVI | 0/0 | 0/0 | Inconclusive |
| Cluster 4 | 367 | 0.358 | 0.356 | 0.244 | Chr. I, II, III, VII, XVI, XX, XXI | 0/0 | 0/0 | Inconclusive |
| Cluster 5 | 47 | 0.463 | 0.494 | 0.027 | Chr. I | 0/0 | 0/0 | Local adaptation (genetic parallelism of **global** freshwater populations) |
| Cluster 6 | 992 | 0.654 | 0.701 | 0.001 | Chr. I (chromosomal inversion) | 32/486 | 4/110 | Local adaptation (genetic parallelism of **global** freshwater populations) |
| Cluster 7 | 32 | 0.575 | 0.597 | 0.094 | Chr. II | 0/0 | 0/0 | Inconclusive |
| Cluster 8 | 34 | 0.331 | 0.354 | NA | Chr. II | 0/0 | 0/0 | Inconclusive |
| Cluster 9 | 39 | 0.301 | 0.290 | 0.803 | Chr. III | 0/0 | 0/0 | Inconclusive |
| Cluster 10 | 70 | 0.467 | 0.497 | 0.002 | Chr. IV | 0/0 | 0/0 | Local adaptation (genetic parallelism of **global** freshwater populations) |
| Cluster 11 | 331 | 0.416 | 0.452 | 0.001 | Chr. IV | 11/166 | 1/13 | Local adaptation (genetic parallelism of **global** freshwater populations) |
| Cluster 12 | 60 | 0.535 | 0.537 | 0.001 | Chr. IV | 3/38 | 0/0 | Local adaptation (genetic parallelism of **global** freshwater populations) |
| Cluster 13 | 115 | 0.702 | 0.715 | 0.027 | Chr. IV | 0/0 | 0/0 | Local adaptation (genetic parallelism of **global** freshwater populations) |
| Cluster 14 | 39 | 0.480 | 0.485 | 1.000 | Chr. IV | 0/0 | 2/4 | Inconclusive |
| Cluster 15 | 54 | 0.498 | 0.527 | 0.393 | Chr. VIII | 0/0 | 0/0 | Inconclusive |
| Cluster 16 | 34 | 0.626 | 0.660 | 0.017 | Chr. IX | 0/0 | 0/0 | Local adaptation (genetic parallelism of **global** freshwater populations) |
| Cluster 17 | 22 | 0.696 | 0.719 | 0.777 | Chr. V | 0/0 | 0/0 | Inconclusive |
| Cluster 18 | 30 | 0.597 | 0.599 | 0.016 | Chr. V | 0/0 | 0/0 | Local adaptation (genetic parallelism of **global** freshwater populations) |
| Cluster 19 | 241 | 0.467 | 0.474 | NA | Chr. V (putative chromosomal inversion) | 0/0 | 0/0 | Putative chromosomal inversion |
| Cluster 20 | 45 | 0.554 | 0.557 | 0.001 | Chr. V | 0/0 | 0/0 | Local adaptation (genetic parallelism of **global** freshwater populations) |
| Cluster 21 | 183 | 0.616 | 0.625 | 0.009 | Chr. X | 0/0 | 0/0 | Local adaptation (genetic parallelism of **Eastern Pacific** freshwater populations) |
| Cluster 22 | 520 | 0.745 | 0.754 | 0.001 | Chr. XI (chromosomal inversion) | 0/0 | 2/4 | Local adaptation (genetic parallelism of **global** freshwater populations) |
| Cluster 23 | 65 | 0.567 | 0.568 | NA | Chr. XIII | 0/0 | 0/0 | Inconclusive |
| Cluster 24 | 47 | 0.581 | 0.586 | 0.006 | Chr. XIV | 0/0 | 0/0 | Local adaptation of two Western Pacific freshwater populations from restricted region |
| Cluster 25 | 36 | 0.455 | 0.475 | 0.035 | Chr. XVI | 0/0 | 0/0 | Local adaptation (genetic parallelism of **global** freshwater populations) |
| Cluster 26 | 71 | 0.587 | 0.601 | NA | Chr. XVI | 0/0 | 0/0 | Inconclusive |
| Cluster 27 | 222 | 0.316 | 0.312 | 0.001 | Chr. XX | 0/0 | 1/87 | Local adaptation (genetic parallelism of **global** freshwater populations) |
| Cluster 28 | 69 | 0.699 | 0.718 | 0.052 | Chr. XX | 0/0 | 0/0 | Inconclusive |
| Cluster 29 | 2730 | 0.691 | 0.704 | 0.001 | Chr. XXI (including a chromosomal inversion) | 0/0 | 30/2551 | Local adaptation (genetic parallelism of **global** freshwater populations but rare outside of EP populations) |

* number of regions/SNPs covered in the top freshwater-marine genomic regions (81 regions) in Jones et al. 2012.

** number of regions/SNPs covered in the 812 regions (after liftover between ref. versions) showing parallel selection in Eastern Pacific freshwater samples identified by SOM/HMM in Jones et al. 2012.
